# Bile acid excess impairs thermogenic function in brown adipose tissue

**DOI:** 10.1101/2020.11.24.396895

**Authors:** Weinan Zhou, Philip VanDuyne, Chi Zhang, Ryan Riessen, Maribel Barragan, Blair M. Rowitz, Margarita Teran-Garcia, Stephen A. Boppart, Sayeepriyadarshini Anakk

## Abstract

Bile acids (BAs) not only facilitate fat digestion but also protect against obesity. Here, we show that a genetic mouse model for BA overload (Farnesoid X receptor; Small heterodimer double knockout (DKO)) exhibits mitochondrial dysfunction resulting in a thermogenic defect. By housing DKO mice at thermoneutrality, the poor mitochondrial function in brown fat protects them from diet-induced obesity. Compared to control, we find higher adipose BA levels with excess accumulation of taurocholic acid in the DKO mice. We report that the expression of genes responsible for BA *de novo* synthesis, conjugation and transporters and accumulation of BAs are present in both brown and white adipocytes. We determine that BA overload is sufficient to cause adipocyte mitochondrial dysfunction and induce the expression of cellular senescence genes *in vitro*. Taken together, we uncover that BA levels within the adipose tissue may modulate its overall function.

**Highlights:**

- Mouse model of BA overload exhibits adipose defects, which is partially restored by housing at thermoneutrality.
- BAs are present in detectable concentrations in both BAT and WAT.
- Adipocytes express genes responsible for *de novo* synthesis, conjugation and transport of BAs, and accumulate BAs.
- Pathological accumulation of BAs impairs mitochondrial function leading to thermogenic defect.

## INTRODUCTION

Adipose tissue is an important regulator of whole-body energy homeostasis. There are two major types of adipose tissues: white (WAT) and brown (BAT). WAT is primarily an energy storage depot, while the energy-burning BAT is responsible for non-shivering thermogenesis (Cohen and Spiegelman, 2016). Adipose tissue undergoes constant remodeling, however, during obesity, excessive expansion occurs resulting in pro-inflammatory milieu and metabolic syndrome (Rosen and Spiegelman, 2014).

Bile acids (BAs) are cholesterol metabolites that are primarily synthesized in the liver and released into the small intestine to facilitate fat digestion and absorption (Chiang, 2004). BAs are also signaling molecules that activate nuclear receptor Farnesoid X receptor (FXR) and membrane G protein-coupled receptor TGR5, to regulate metabolic homeostasis in the liver, intestine and adipose tissue (Chavez-Talavera et al., 2017; Li and Chiang, 2014; Yang et al., 2018). Earlier studies have shown that physiological levels of circulating BAs can promote heat production in the BAT to increase energy expenditure and prevent diet-induced obesity via the TGR5 signaling pathway (Broeders et al., 2015; Watanabe et al., 2006). However, the BA concentrations and their role within the adipose tissue has not been fully explored. We examined and found that the mouse model of BA overload displayed a thermogenic defect. We tested if the defective thermogenesis is due to BAs. Then, we examined the concentrations and composition of BAs in brown and white fat depots. We also investigated whether genes involved in BA synthesis and transport are expressed *de novo* in the adipose tissue, and if BAs are accumulated within adipocytes. Finally, we evaluated the mitochondrial function and cellular senescence of adipocytes in the presence and absence of BAs *in vitro*. Intriguingly, our results indicate that the defective thermogenesis subsequent to excess BAs may contribute towards protection against diet-induced obesity in mice.

## RESULTS

### DKO Mice Display Reduced Fat Accumulation and Impaired Brown Adipose Mitochondrial Function

Analysis of the genetic mouse model of nuclear receptors (Farnesoid X receptor (FXR), Small heterodimer partner(SHP)) double knockout (DKO) mice revealed elevated levels of hepatic (Table 1) and circulating BAs as we previously described (Anakk et al., 2011; Desai et al., 2017).These DKO mice show lower body weight and resistance to fatty liver disease (Akinrotimi et al., 2017). These mice also exhibit decreased brown and white fat mass (Figure 1A) and reduced adipocyte size (Figures 1B, 1C and S1A) compared to WT mice even under normal chow diet. This finding correlated well with reduced expression of lipogenic genes in the BAT (*C/Ebpa, Srebp1c*) and WAT (*C/Ebpa, Fasn*) (Figure S1B), suggesting reduced lipid synthesis in DKO adipose tissue. Although, DKO WAT displayed increased mRNA levels of SCD1 (Figure S1B), a rate-limiting enzyme for the synthesis of unsaturated fatty acids (Lodhi et al., 2011), no alteration was seen in the degrees of unsaturation in both WAT and BAT compared to WT mice (Figure S1D).

**Table 1.**
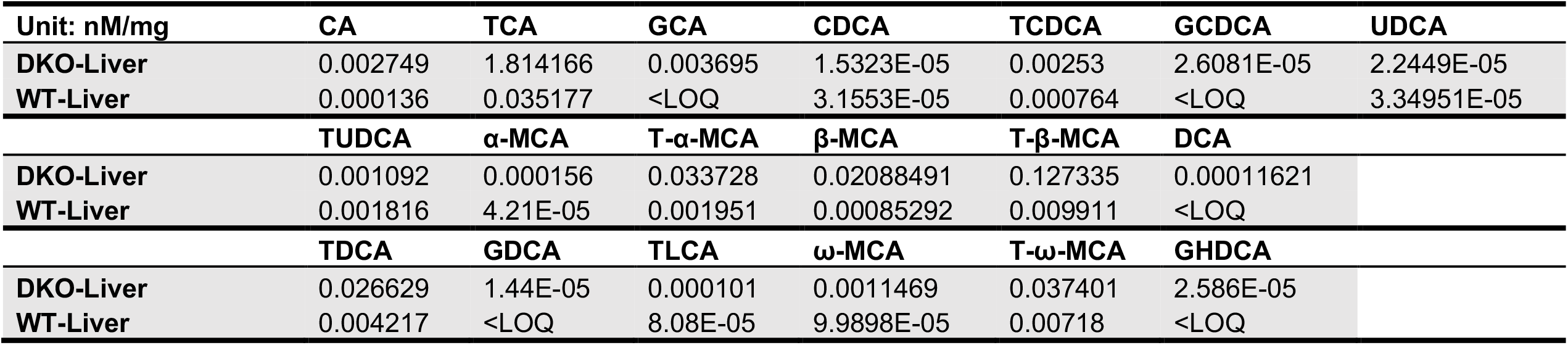
Hepatic bile acid concentrations in DKO and WT mice.

| Unit: nM/mg | CA | TCA | GCA | CDCA | TCDCA | GCDCA | UDCA |
| --- | --- | --- | --- | --- | --- | --- | --- |
| DKO-Liver | 0.002749 | 1.814166 | 0.003695 | 1.5323E-05 | 0.00253 | 2.6081E-05 | 2.2449E-05 |
| WT-Liver | 0.000136 | 0.035177 | <LOQ | 3.1553E-05 | 0.000764 | <LOQ | 3.34951E-05 |
| | TUDCA | α-MCA | T-α-MCA | β-MCA | T-β-MCA | DCA | |
| DKO-Liver | 0.001092 | 0.000156 | 0.033728 | 0.02088491 | 0.127335 | 0.00011621 |  |
| WT-Liver | 0.001816 | 4.21E-05 | 0.001951 | 0.00085292 | 0.009911 | <LOQ |  |
| | TDCA | GDCA | TLCA | ω-MCA | T-ω-MCA | GHDCA | |
| DKO-Liver | 0.026629 | 1.44E-05 | 0.000101 | 0.0011469 | 0.037401 | 2.586E-05 |  |
| WT-Liver | 0.004217 | <LOQ | 8.08E-05 | 9.9898E-05 | 0.00718 | <LOQ |  |

**Figure 1.**
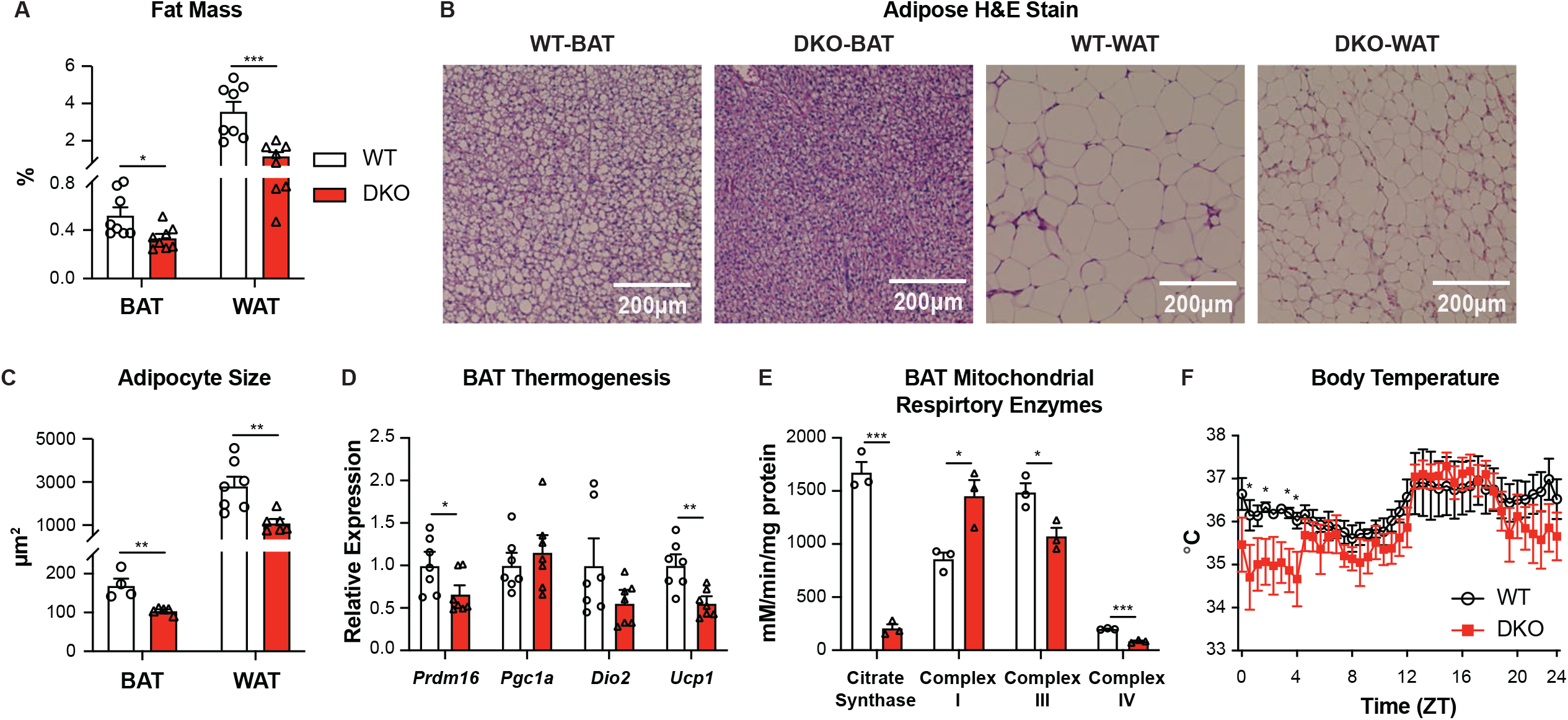
DKO mice show decreased fat accumulation and BAT mitochondrial dysfunction upon normal chow diet. (A) Brown and white fat mass/body weight ratios of DKO and WT mice upon normal chow (n=8 per group). (B-C) Representative images (B) (scale bar: 200 μm) and quantification of adipocyte size (C) of H&E-stained BAT (n=4-5 mice per group) and WAT (n=6-7 mice per group) sections of DKO and WT mice upon normal chow. (D-E) Thermogenic gene expression (D) (n=7 mice per group) and mitochondrial respiratory enzyme activity (E) (n=3 mice per group) of the BAT from WT and DKO mice upon normal chow. (F) Body temperature of DKO and WT mice for 24 hours upon normal chow (n=7 mice per group). Data are represented as mean ± SEM. \**P* < 0.05, \*\**P* < 0.01, \*\*\**P* < 0.001 compared to WT mice.

Since BAT mitochondrial thermogenesis can promote fat burning (Cohen and Spiegelman, 2016), we examined mitochondrial function in the BAT and continually monitored the body temperature of DKO and WT mice. Surprisingly, DKO mice exhibited decreased expression of thermogenic genes *Prdm16* and *Ucp1*, while other mitochondrial genes *Pgc1a* and *Dio2* were unchanged (Figure 1D). Despite an increase in respiratory chain complex I activity, we determined significantly reduced activities of citrate synthase as well as respiratory chain complex III and IV (Figure 1E). Further, DKO mice exhibited lower body temperature than WT mice during the daytime but not the night (Figure 1F). WAT, on the other hand, displayed comparable levels of mitochondrial genes between DKO and WT mice when housed at RT (Figure S2A). These results suggest that DKO mice have decreased lipid biosynthesis, poor mitochondrial function and impaired heat production in the brown fat.

### Thermoneutral Housing Negates Brown Fat Defect and Restore Weight Gain upon High-fat Diet Condition in the DKO mice

To confirm impaired brown adipose mitochondrial function, we housed both WT and DKO mice at thermoneutrality (TN, 30 °C), which excludes the BAT thermogenic effect (Boutant et al., 2017). TN reduced the expression of crucial thermogenic genes, *Dio2* and *Ucp1*, in the BAT from both WT and DKO mice (Figure S2B). However, TN neither altered mRNA levels of thermogenic transcription factors *Prdm16* and *Pgc1a* in the BAT nor WAT mitochondrial genes including *Prdm16, Pgc1a, Dio2* and *Cpt1a* in WT mice (Figures S2A and S2B). Despite a slightly smaller adipocyte size, the brown fat mass was comparable between DKO and WT BAT (Figures 2A-2C and S3A). Similarly, the transcript profile of lipogenic genes *C/Ebpa* and *Srebp1c* between DKO and WT mice housed at 30 °C was not different (Figure S3B). However, TN did not alter the reduction in white fat mass (Figure 2A) and adipocyte size (Figures 2B-2C and S3A), or the induction in WAT lipogenic gene *Scd1* expression (Figure S3B) in DKO mice. These results indicate poor brown fat function may protect the DKO mice against obesity. If this is true, then the DKO mice housed at 30 °C will lose their protection and gain weight.

**Figure 2.**
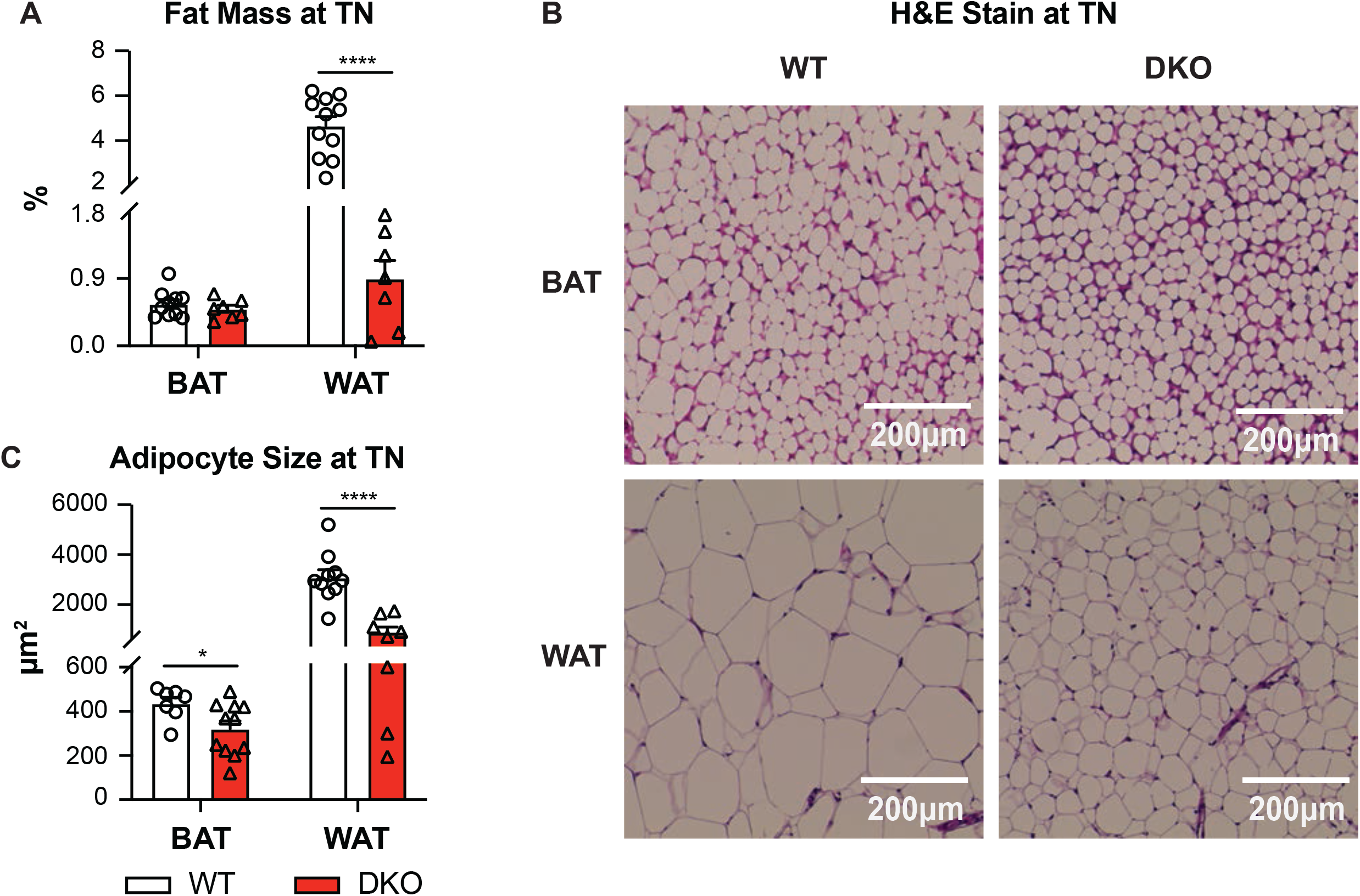
Thermoneutrality abolishes DKO BAT defect in fat accumulation upon normal chow diet. (A) Brown and white fat mass/body weight ratios of DKO and WT mice upon normal chow housed at TN for 8 weeks (n=7-11 per group). (B-C) Representative images (B) (scale bar: 200 μm) and quantification of adipocyte size (C) of H&E-stained BAT (n=7-11 mice per group) and WAT (n=8-10 mice per group) sections of DKO and WT mice upon normal chow housed at TN. Data are represented as mean ± SEM. \**P* < 0.05, \*\*\*\**P* < 0.0001 compared to WT mice.

We then challenged DKO and WT mice with a 45% high-fat diet (HFD) for 8 weeks at either RT or TN conditions. DKO mice gained a similar percentage in body weight as WT (Figure 3A) in response to HFD at TN and showed increased BAT mass but not in WAT mass (Figure 3B). Although at RT, the adipocytes were smaller in both brown and white fat from DKO than WT mice upon HFD (Figures 3C, 3D and S4A), TN housing led to an increase in BAT adipocyte size in both WT and DKO mice (Figures 3C, 3D and S4A). Housing did not alter the WAT adipocyte size (Figures 3C, 3D and S4A). Similarly, WAT gene expression profile showed that lipogenic genes were comparable between DKO and WT mice at RT (Figure S4B).

**Figure 3.**
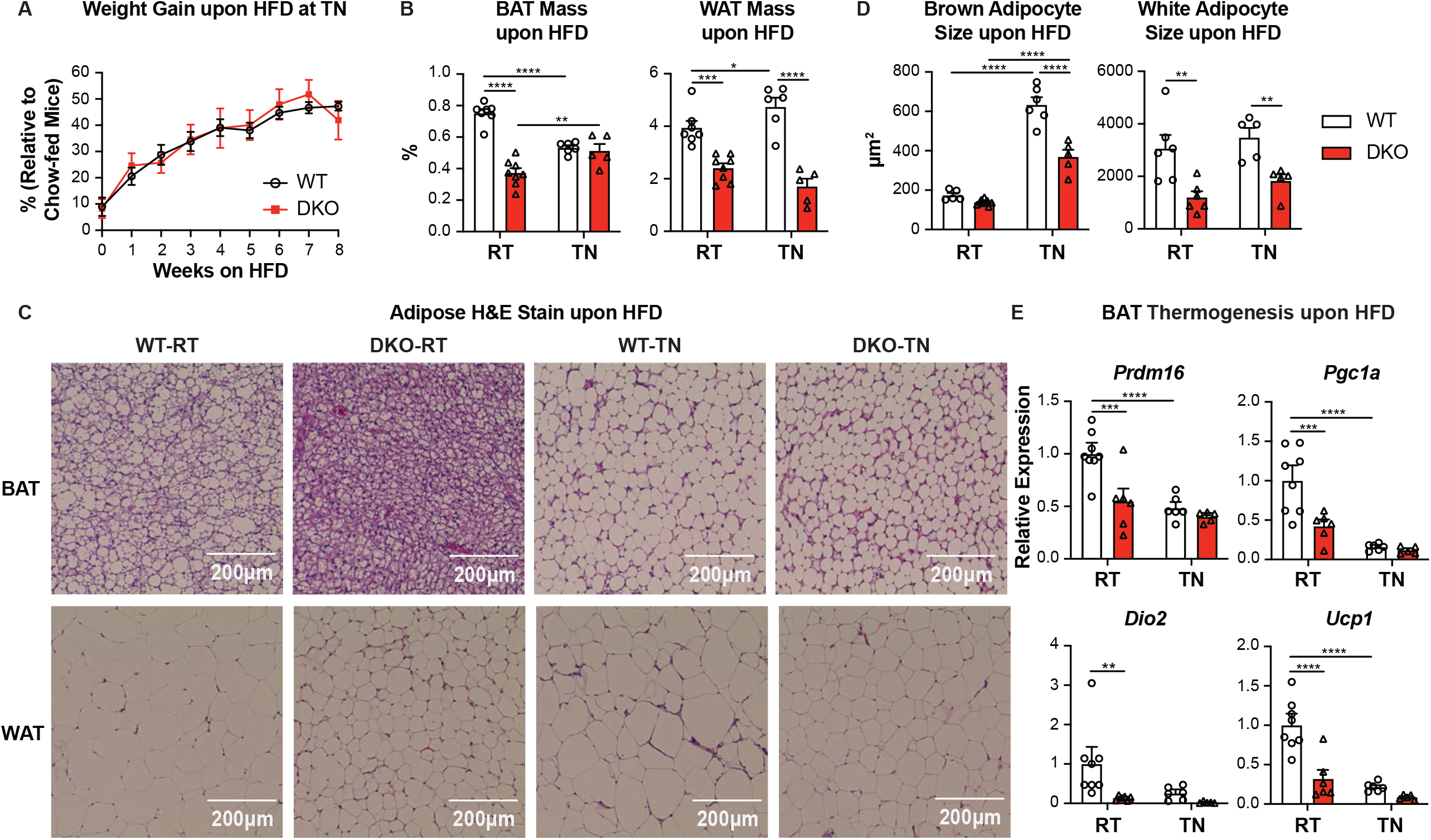
DKO mice display reduced fat accumulation upon HFD. (A-B) Body weight gain (A) (n=6-7 mice per group) and fat mass/body weight ratios (B) (n=7-8 mice per group for RT, n=5-6 mice per group for TN) of DKO and WT mice housed at RT or TN upon HFD. (C-D) Representative images (C) (scale bar: 200 μm) and quantification of adipocyte size (D) of H&E-stained BAT (n=5-9 mice per group for RT, n=5-6 mice per group for TN) and WAT (n=6 mice per group for RT, n=5 mice per group for TN) sections from WT and DKO mice housed at RT or TN upon HFD. (E) Expression of thermogenic genes of BAT from DKO and WT mice housed at RT (n=6-8 mice per group) or TN (n=5-6 mice per group) upon HFD for 8 weeks. Data are represented as mean ± SEM. \**P* < 0.05, \*\**P* < 0.01, \*\*\**P* < 0.001, \*\*\*\**P* < 0.0001.

However, WT BAT at TN post-HFD revealed significant induction of *Srebp1c* while *Pparg, Fasn* and *Scd1* were suppressed with no change in *C/Ebpa* transcript levels (Figure S4C). Despite lower expression of many of these genes including *C/Ebpa* levels at RT, DKO BAT was able to upregulate *C/Ebpa* as well as *Srebp1c* transcript levels to that of WT at 30°C (Figure S4C), while *Pparg, Fasn* and *Scd1* were maintained at the reduced levels (Figure S4C). These results indicate that fat accumulation in the BAT of DKO mice under TN correlates to increasing *C/EBPa* and *Srebp1c* levels.

Similar to the findings upon chow, HFD-fed DKO mice also showed lower expression of thermogenic genes *Prdm16, Pgc1a, Dio2* and *Ucp1* in the BAT compared to HFD fed WT mice when housed at RT (Figure 3E). Moreover, levels of BAT thermogenic genes including *Prdm16, Pgc1a* and *Ucp1* were suppressed in WT mice upon housing at 30°C to that of DKO BAT at TN (Figure 3E). On the other hand, housing conditions modestly altered WAT gene expression pattern except for *Prdm16* expression, which was higher in the DKO mice only at RT. *Pgc1a* was induced, while *Dio2* and *Cpt1a* levels were reduced in DKO compared to WT WAT at both housing conditions after HFD (Figure S5). Next, we examined whether the compromised function is accompanied with abnormal structure of mitochondria. Under 30°C housing, the total number of mitochondria including the ones with abnormal structural features increased, whereas mitochondria with the normal structure were reduced compared to the mice housed at RT (Figures 4A-4C). These structural changes were seen in both the genotypes. But, HFD fed DKO mice housed at TN displayed an exaggerated mitochondrial change compared to HFD fed WT mice. The ratio of normal mitochondria in the DKO BAT dramatically dropped with the majority of them revealing abnormal ultrastructure including irregular shape, loss of cristae or presence of myelin figures (Figures 4A and 4C), an indicator of mitochondrial damage or degeneration (Le Beux et al., 1969; Siskova et al., 2010).

**Figure 4.**
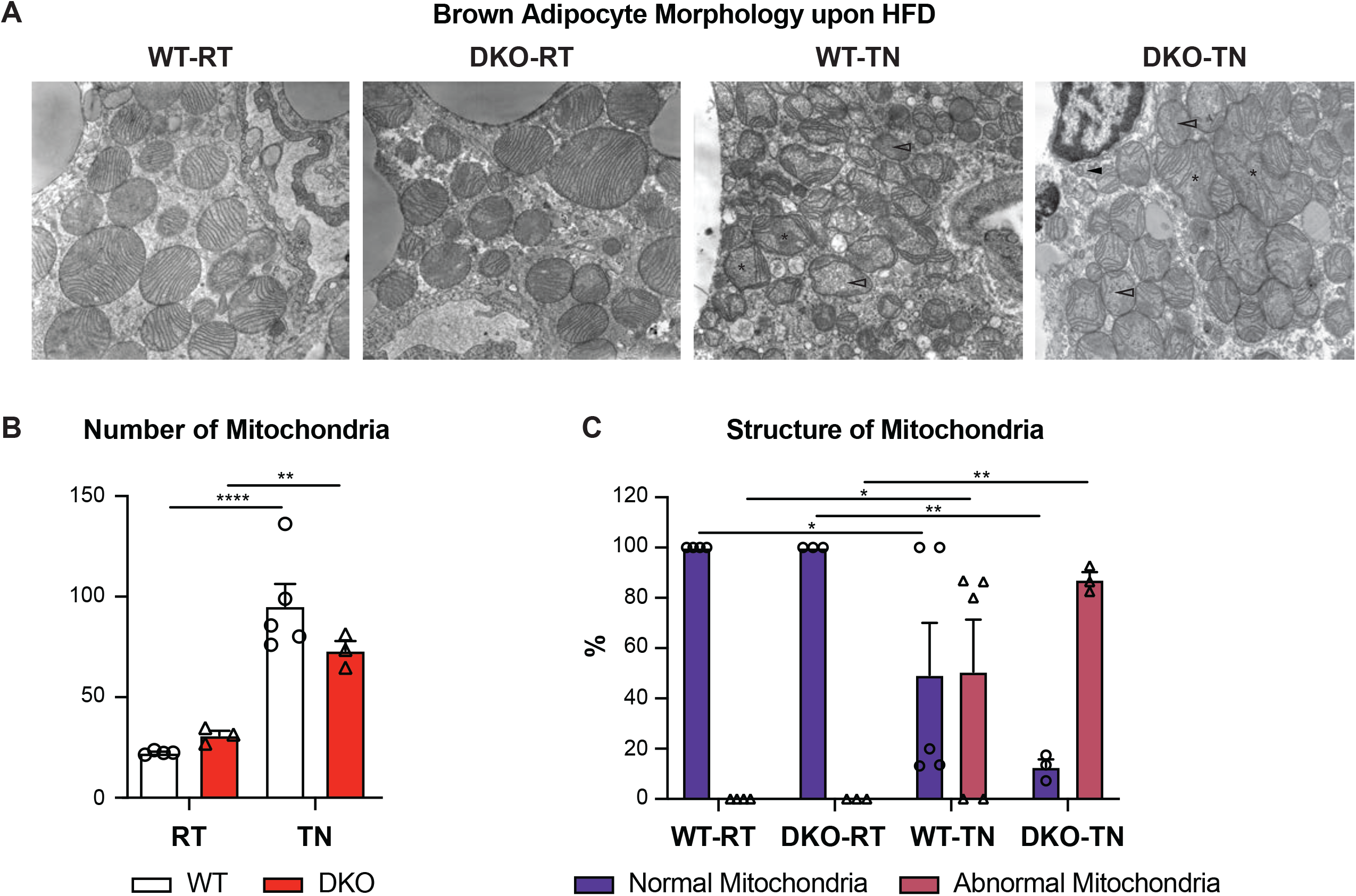
Thermoneutral housing alters BAT mitochondrial number and structure upon HFD. (A-C) Representative electron microscopy images (A) (abnormal mitochondria with irregular shape (asterisks), loss of cristae (empty arrowheads) or myelin figures (arrowheads), magnification: 10,000x) as well as quantification of mitochondria number (B) and structure (C) of BAT from DKO and WT mice housed at RT (n=3-4 mice per group) or TN (n=3-5 mice per group) upon HFD for 8 weeks. Data are represented as mean ± SEM. \**P* < 0.05, \*\**P* < 0.01, \*\*\*\**P* < 0.0001.

### BAs are Present in the Adipose Tissue and Their Composition is Altered in the DKO Mice

BAs have been shown to regulate lipid metabolism (Chavez-Talavera et al., 2017; Li and Chiang, 2014; Yang et al., 2018), so we investigated the local levels of BAs in the fat depots. Previous studies have reported the presence of BAs in the white fat of humans (Jantti et al., 2014) and mice (La Frano et al., 2017). We analyzed and found detectable levels of BAs in both BAT and WAT. BA concentrations (Figure 5A and Table 2), composition (Figure 5B) and hydrophobicity indices (Figure 5C) were comparable between the two adipose depots of WT mice. However, DKO mice displayed higher concentrations of BAs in both depots (~16x more in the BAT and ~6x in the WAT) and also revealed composition differences compared to the WT mice (Figures 5A and 5B). Both adipose depots in the DKO mice showed about a 3-fold increase in taurocholic acid (TCA) with a concomitant decrease in cholic acid (CA) indicating increased BA-conjugation. We also found an increase in β-muricholic acid (β-MCA) in the brown fat of DKO mice. Intriguingly, DKO BAT showed reductions in secondary BAs that are generated by dehydroxylation or epimerization by the gut microbiota. For instance, deoxycholic acid (DCA) and tauro-ω-muricholic acid (T-ω-MCA) were decreased compared to WT mice. Similarly, DKO WAT also showed a robust decrease in secondary BA, DCA, but pronounced increases in primary BA, tauro-α-muricholic acid (T-α-MCA) (Figure 5B). Albeit high concentrations in the BAT and WAT, the composition changes result in a decrease in BA hydrophobicity in DKO compared to WT mice (Figure 5C).

**Table 2.** Adipose bile acid concentrations in DKO and WT mice.

| Unit: nM/mg | CA | TCA | GCA | CDCA | TCDCA | GCDCA | UDCA | TUDCA |
| --- | --- | --- | --- | --- | --- | --- | --- | --- |
| DKO-BAT-1 | 0.006076 | 0.043023 | 0.000147 | 1.29E-05 | 5.53E-05 | 2.56E-06 | 8.88E-05 | 0.000136 |
| DKO-BAT-2 | 0.003566 | 0.00597 | 2.56E-06 | 3.02E-05 | 2.56E-06 | 1.18E-05 | 2.37E-05 | 2.56E-06 |
| DKO-BAT-3 | 0.002302 | 0.006983 | 2.15E-05 | 2.97E-05 | 2.56E-06 | 2.56E-06 | 1.57E-05 | 2.24E-05 |
| WT-BAT-1 | 0.000454 | 0.00017 | 2.56E-06 | 1.45E-05 | 2.56E-06 | 2.56E-06 | 2.56E-06 | 2.56E-06 |
| WT-BAT-2 | 0.000558 | 0.000191 | 2.56E-06 | 1.06E-05 | 2.56E-06 | 2.56E-06 | 2.61E-05 | 2.56E-06 |
| WT-BAT-3 | 0.002769 | 0.000236 | 2.56E-06 | 2.56E-06 | 2.56E-06 | 2.56E-06 | 2.56E-06 | 2.56E-06 |
| DKO-Gonadal WAT-1 | 0.000901 | 0.019745 | 6.78E-05 | 2.56E-06 | 4.94E-05 | 2.56E-06 | 2.14E-05 | 7.55E-05 |
| DKO-Gonadal WAT-2 | 0.002032 | 0.001862 | 2.56E-06 | 5.87E-05 | 2.56E-06 | 2.56E-06 | 4.49E-05 | 2.56E-06 |
| DKO-Gonadal WAT-3 | 0.002705 | 0.002086 | 2.56E-06 | 0.000057 | 2.56E-06 | 2.56E-06 | 0.000072 | 2.56E-06 |
| WT-Gonadal WAT-1 | 0.000361 | 0.000102 | 2.56E-06 | 7.98E-06 | 0 | 2.56E-06 | 2.56E-06 | 2.56E-06 |
| WT-Gonadal WAT-2 | 0.002127 | 0.00071 | 2.56E-06 | 9.17E-05 | 2.56E-06 | 1.23E-05 | 3.01E-05 | 2.56E-06 |
| WT-Gonadal WAT-3 | 0.001886 | 0.000152 | 2.56E-06 | 2.56E-06 | 2.56E-06 | 2.56E-06 | 2.56E-06 | 2.56E-06 |
| | α-MCA | T-α-MCA | β-MCA | T-β-MCA | DCA | TDCA | ω-MCA | T-ω-MCA |
| DKO-BAT-1 | 2.65E-05 | 0.002718 | 0.001613 | 0.003223 | 0.001776 | 0.000955 | 0.000293 | 0.001477 |
| DKO-BAT-2 | 2.56E-06 | 0.000252 | 0.000204 | 0.000284 | 0.000274 | 0.000131 | 7.27E-05 | 0.000141 |
| DKO-BAT-3 | 0 | 0.00013 | 0.000282 | 0.00049 | 0.000141 | 0.000358 | 2.56E-06 | 0.000143 |
| WT-BAT-1 | 2.56E-06 | 0.000029 | 2.56E-06 | 7.52E-05 | 0.000142 | 2.24E-05 | 0 | 4.25E-05 |
| WT-BAT-2 | 2.56E-06 | 2.74E-05 | 2.56E-06 | 0.000062 | 8.66E-05 | 2.31E-05 | 2.56E-06 | 6.13E-05 |
| WT-BAT-3 | 2.56E-06 | 2.56E-06 | 2.56E-06 | 4.39E-05 | 5.07E-05 | 2.64E-05 | 2.56E-06 | 4.24E-05 |
| DKO-Gonadal WAT-1 | 2.56E-06 | 0.001603 | 0.000987 | 0.002692 | 0.000619 | 0.000499 | 0.000235 | 0.000994 |
| DKO-Gonadal WAT-2 | 2.56E-06 | 0.000111 | 4.46E-05 | 0.000148 | 9.64E-05 | 0.000057 | 2.56E-06 | 8.16E-05 |
| DKO-Gonadal WAT-3 | 2.56E-06 | 6.94E-05 | 5.78E-05 | 0.000166 | 6.72E-05 | 4.56E-05 | 2.56E-06 | 4.95E-05 |
| WT-Gonadal WAT-1 | 2.56E-06 | 2.56E-06 | 2.56E-06 | 3.43E-05 | 1.85E-05 | 2.56E-06 | 2.56E-06 | 1.95E-05 |
| WT-Gonadal WAT-2 | 2.56E-06 | 0.000105 | 2.56E-06 | 0.000276 | 0.000429 | 5.24E-05 | 2.56E-06 | 0.00013 |
| WT-Gonadal WAT-3 | 2.56E-06 | 2.74E-05 | 2.56E-06 | 9.47E-05 | 2.56E-05 | 2.56E-06 | 0 | 6.14E-05 |

**Figure 5.**
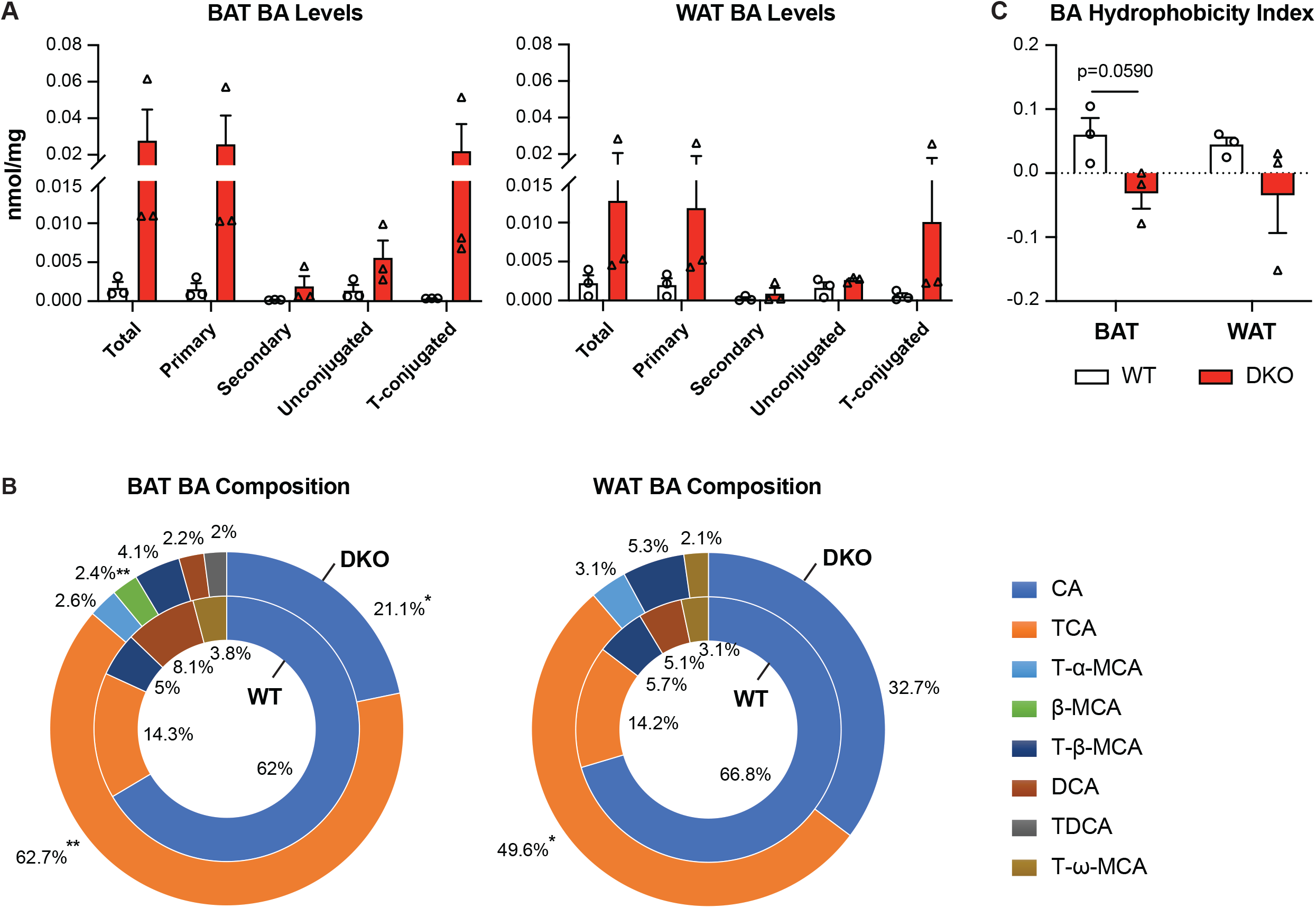
DKO mice exhibit altered adipose BA composition. (A-C) Levels (A), composition (B) (BA percentage below 2% is not shown) and hydrophobicity (C) of BAs in the BAT and WAT from DKO and WT mice upon normal chow (n=3 mice per group). Data are represented as mean ± SEM. \**P* < 0.05, \*\**P* < 0.01 compared to WT mice.

### Adipocytes Express Genes Regulating the Synthesis, Conjugation and Transport of BAs and Accumulate BAs

The elevated concentrations of primary BAs in the adipose tissues raise the question if they can be synthesized or transported locally. To address this, we examined (i) the expression of genes controlling BAs synthesis and transport from the systemic circulation into cells within the adipose tissue, as well as (ii) BA accumulation within adipocytes. We found the expression of BA synthesis genes in both mouse and human adipose tissues (Figures 6A and 6B). Brown fat of WT mice expressed higher levels of BA synthesis genes *Cyp7a1, Cyp27a1* and *Cyp2c70* but lower levels of *Cyp7b1* compared to the white fat (Figure 6A). Human WAT also expressed these BA synthesis genes with *CYP8B1* being the highest expressed (Figure 6B). To detect if primary adipocytes reveal the same expression pattern, we investigated pre- and post-differentiated primary adipocytes derived from either brown or white adipose depots. Primary adipocytes from both depots displayed expression of all the BA synthesis genes with higher *Cyp7b1* and *Cyp8b1* levels in the preadipocytes derived from the WAT compared to BAT. Both BAT and WAT preadipocytes exhibited a minimal *Cyp27a1* and no detectable *Cyp2c70* expression which were dramatically induced post-differentiation in these cultured cells. Further, *Cyp8b1* levels were upregulated in the differentiated brown adipocyte cultures while unaltered in the white adipocyte cultures. *Cyp7b1* levels were reduced after differentiation in both white and brown adipocyte cultures, which matched with the tissue expression pattern. *Cyp7a1* expression remained unaltered irrespective of the origin of the preadipocyte culture and its differentiated state (Figures 6C). Next, we examined and found induction of BA conjugation genes *Slc27a5* and *Baat* as well as transporter genes *Slco1a6, Slco1b2* and *Slc51b* in differentiated brown adipocyte cultures (Figures 6D and 6E). Conversely, in the adipocytes derived from white adipose, we found that the expression of genes regulating BA conjugation and transport including *Baat, Slco1a6, Slco1b2, Slc51a* and *Slc51b* remained lower than the brown adipocytes and were unaltered irrespective of their differentiation status (Figures 6D and 6E). Further, we found the accumulation of fluorescent BAs, choly-lysyl-fluorescein (CLF), within both brown and white differentiated primary adipocytes upon CLF treatment (Figure 7). These findings demonstrate that genes regulating BA synthesis, conjugation and transport are expressed in adipose depots and they exhibit distinct expression profile between brown and white adipocytes. Importantly, functional BA uptake may occur in adipocytes.

**Figure 6.**
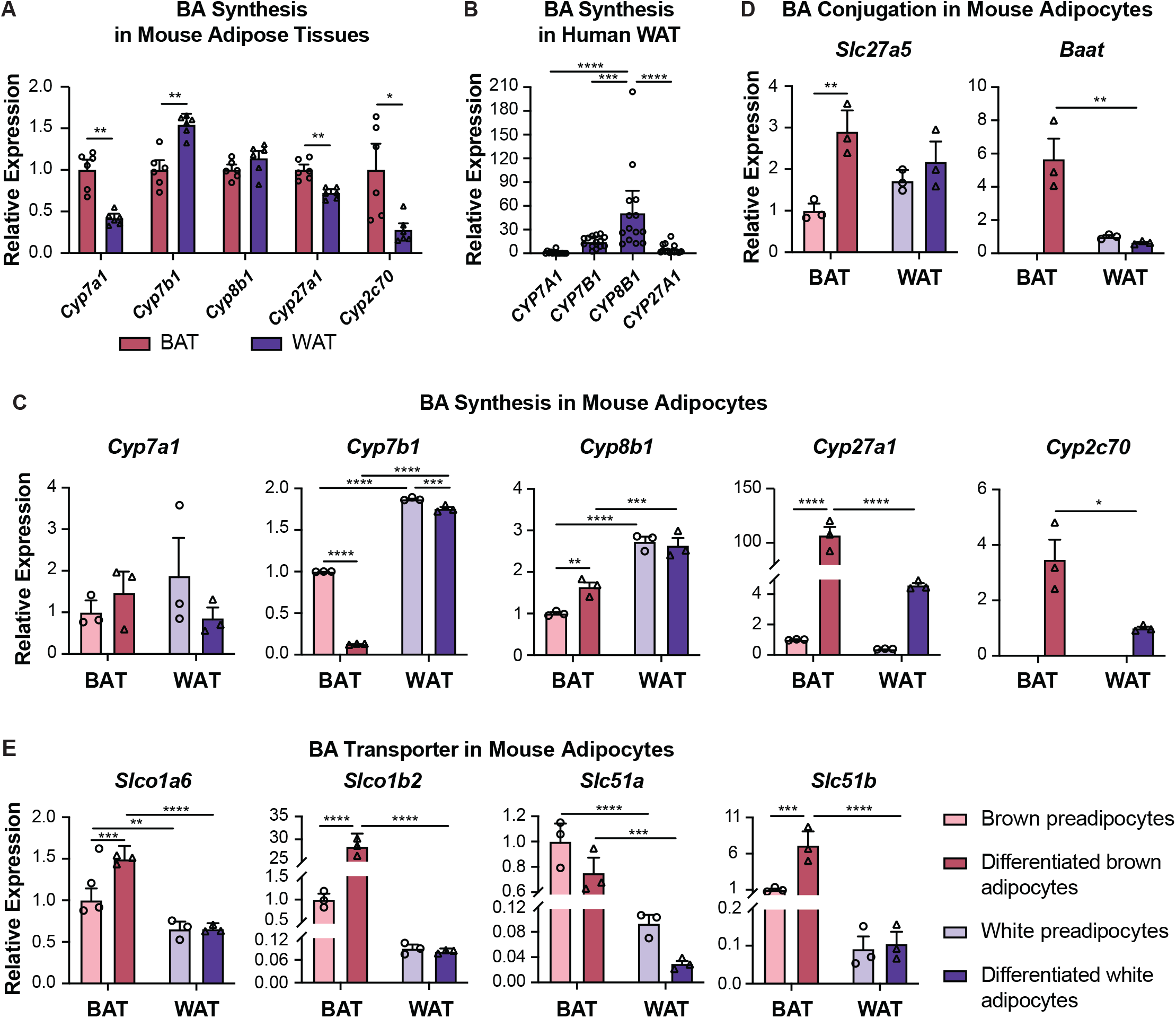
BA synthesis, conjugation and transporter genes are expressed in adipocytes. A-B) Expression of BA synthesis genes in the BAT and WAT from chow-fed WT mice (A) (n=6 mice) and in the perigastric WAT from obese humans (B) (n=14 individuals). (C-E) Expression of BA synthesis (C), conjugation (D) and transporter (E) genes in primary pre- and post-differentiated adipocytes from BAT and WAT of chow-fed WT mice (n=3 cultures). Data are represented as mean ± SEM. \**P* < 0.05, \*\**P* < 0.01, \*\*\**P* < 0.001, \*\*\*\**P* < 0.0001.

**Figure 7.**
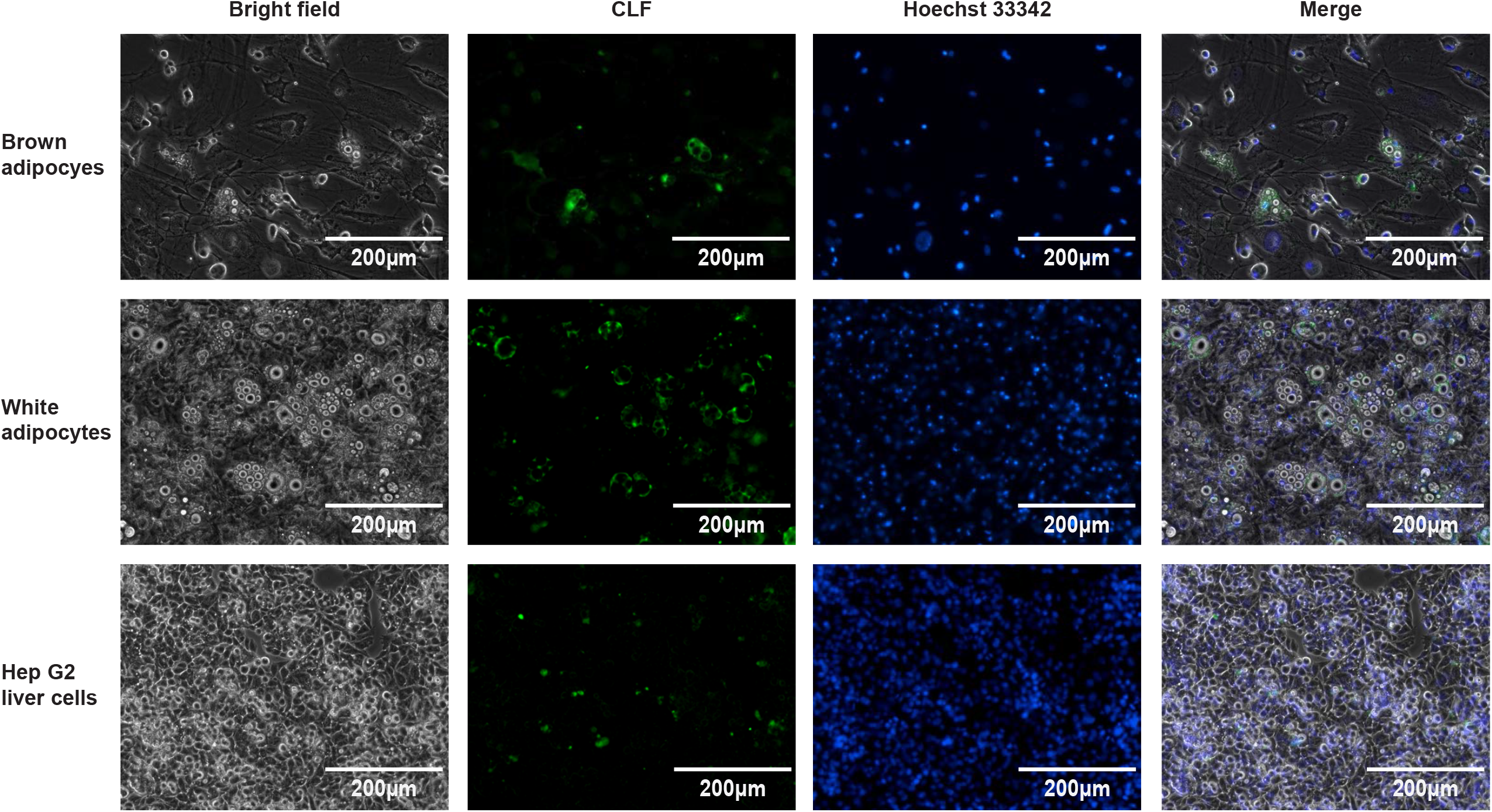
BAs are accumulated within adipocytes. Representative images of accumulation of a fluorescein-labeled BA, CLF, within the differentiated brown and white adipocytes as well as Hep G2 liver cell line (positive control) upon CLF treatment for 24 hours (scale bar: 200 μm; n=3 cultures per group). Nuclei were stained with Hoechst 33342.

### High BA Levels Reduce Mitochondrial Membrane Potential and Induce Cellular Senescence Gene Expression in Adipocytes

Based on our findings that DKO BAT exhibit mitochondrial defect and that they accumulate high BA concentrations, we examined the role of BAs in adipocyte mitochondrial function. We treated differentiated primary adipocytes derived from either BAT or WAT with pathological concentrations of chenodeoxycholic acid (CDCA). We observed that high doses of CDCA did not affect cell viability (Figure S6), but was sufficient to reduce mitochondrial membrane potential (Figures 8A and 8B) in both brown and white adipocytes. Further, CDCA treatment significantly decreased the expression of thermogenic genes including *Prdm16, Pgc1a, Dio2* and *Ucp1* in the adipocytes derived from brown fat (Figures 8C). Except for *Prdm16, Pgc1a, Dio2* and *Cpt1a* levels were reduced upon CDCA treatment in white adipocytes as well (Figure 8D). These results reveal that the accumulation of CDCA can dramatically reduce mitochondrial membrane potential and gene expression in adipocytes.

**Figure 8.**
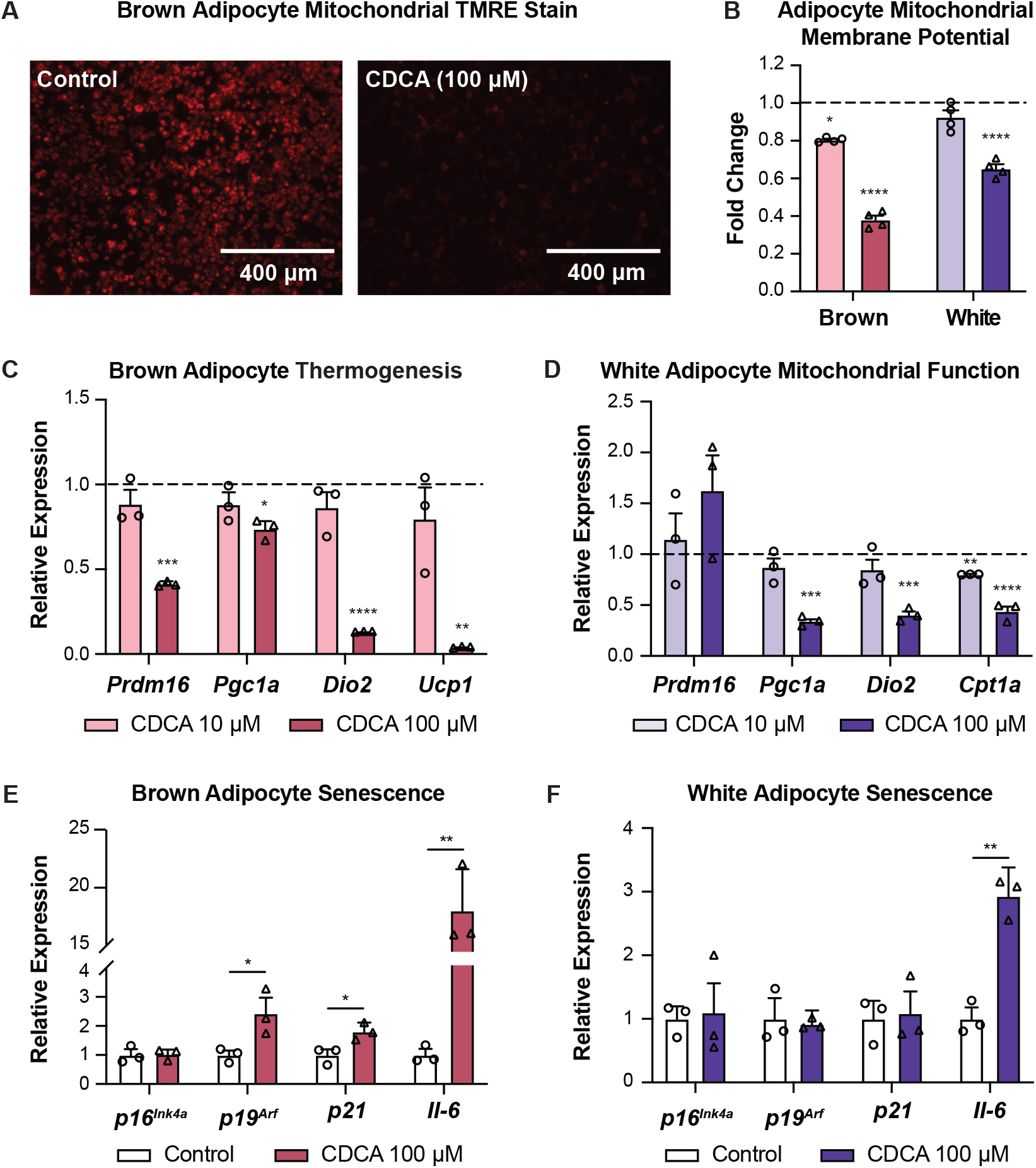
Pathological concentrations of CDCA impair adipocyte mitochondrial function and induce senescence gene expression. (A-B) Representative images of TMRE staining of active mitochondria (A) (scale bar: 400 μm) and quantification of mitochondrial membrane potential (B) of differentiated brown and white adipocytes upon CDCA treatment for 24 hours (n=4 cultures per group). (C-D) Expression of genes involved in thermogenesis in differentiated brown adipocytes (C) as well as genes regulating mitochondrial function in differentiated white adipocytes (D) in response to pathological concentrations of CDCA (n=3 cultures per group). (E-F) Expression of senescence marker genes in differentiated brown (E) and white (F) adipocytes in response to pathological concentrations of CDCA (n=3 cultures per group). Data are represented as mean ± SEM. \**P* < 0.05, \*\**P* < 0.01, \*\*\**P* < 0.001, \*\*\*\**P* < 0.0001 compared to vehicle-treated adipocytes indicated by the dashed line.

Aging is a risk factor for disease and death (Harman, 1991), and is associated with fat loss, mitochondrial dysfunction and impaired thermogenesis in the BAT (Florez-Duquet and McDonald, 1998) as well as increased BA levels (Lee et al., 2016). To explore whether high BAs accelerate the aging of adipose tissue, we examined cellular senescence, the critical process underlying aging, in response to CDCA treatment, and found that high levels of CDCA increased the expression of senescence genes including *p19*^*Arf*^, *p21* and *Il-6* in brown adipocytes (Figure 8E). CDCA treatment upregulated *Il-6* without affecting the levels of *p16*^*Ink4a*^, *p19*^*Arf*^ and *p21* in white adipocytes (Figure 8F). These findings demonstrate that excessive CDCA may induce brown adipocyte senescence.

## DISCUSSION

Several studies have shown that BAs can alleviate diet-induced obesity through activating brown fat (Broeders et al., 2015; Watanabe et al., 2006). While BA role in the regulation of lipid metabolism has been studied (Chavez-Talavera et al., 2017; Li and Chiang, 2014; Yang et al., 2018), their *de novo* function within the adipocytes is poorly understood.

We used the global *Fxr*; *Shp* DKO mouse model for BA overload (Anakk et al., 2011; Desai et al., 2017) that shows resistance to diet-induced obesity (Akinrotimi et al., 2017). To explore the adipose function, we examined BA concentrations and composition in both brown and white fat and found them elevated in DKO compared to WT mice. The systemic circulation can contribute to adipose BA levels (Jantti et al., 2014), therefore, we tested BA uptake transporters and found *Slco1a6* (Tian et al., 2015), *Slco1b2* (Csanaky et al., 2011; Slijepcevic et al., 2017) and *Slc51a/b* (Suga et al., 2019) were expressed in the adipose tissue. Surprisingly, we visualized the accumulation of a conjugated-cholic acid (CA) analogue by adipocytes. Due to the hydrophilic property of conjugated-CA (Roda et al., 1990), these results suggest that BAs may be taken up by adipocytes using a transporter-medicated mechanism. On the other hand, BA conjugation genes were robustly expressed in adipocytes indicating the possibility of *de novo* taurine conjugation within the adipose. Importantly, DKO mice showed decreased BA hydrophobicity in the adipose but increased BA hydrophobicity in the serum (Anakk et al., 2011). In particular, the levels of hydrophilic BAs α/β-MCA significantly increased in the fat depots while only moderately upregulated in the serum of DKO compared to WT mice (Desai et al., 2017). These results imply that *de novo* synthesis may contribute to the hydrophilic BAs in the adipose tissue. Taken together, these data suggest that both BA *de novo* synthesis and transport may contribute to the differential adipose BA composition and concentrations in DKO mice. Further studies will be needed to characterize functional BA transporters and the *de novo* synthesis pathway. Our findings are consistent with a recent report that shows the presence of BAs in the BAT (Zhang et al., 2019). To overcome the caveat of whole-body knockout of *Fxr* and *Shp*, we performed *in vitro* studies using primary adipocyte cultures with intact FXR and SHP signaling to tease apart the BA effect. Previous studies have shown that *Fxr* or *Shp* deletion does not alter the basal BAT thermogenic gene expression profile (Park et al., 2011; Zhang et al., 2012).

Since we observed mitochondrial defects in the BAT of DKO mice, we investigated if high concentrations of BAs are detrimental and affect adipocyte mitochondrial function. In line with previous findings that physiological levels of BAs promote brown fat activity (Broeders et al., 2015; Watanabe et al., 2006), low concentrations of CDCA increase the mitochondrial membrane potential of brown adipocytes (Figure S7). However, when the BA concentration exceeds 100 μM as seen in the DKO mice, it impairs mitochondrial function (Figure S7 and 8C). Due to their detergent properties, intracellular accumulation of BAs can increase mitochondrial membrane permeability leading to mitochondrial depolarization, which eventually triggers cell injury or death in hepatocytes (Rodrigues and Steer, 2000; Rolo et al., 2000; Sokol et al., 2005) and cardiomyocytes (Ferreira et al., 2005). Interestingly, despite weakening mitochondrial function, elevated doses of BAs did not cause any defects in adipocyte viability. Because of hydrophobic BA-induced cell injury and hydrophilic BA-induced cell protection (de Aguiar Vallim et al., 2013; Katagiri et al., 1992; Perez and Briz, 2009), the cytotoxicity of high hydrophobic BAs may be ameliorated by the altered BA composition. Thus, both elevated levels and altered composition of BAs may modulate adipose mitochondrial function. Importantly, we detected that the poor mitochondrial function in brown fat can partially protect the DKO mice against obesity. Previous studies have reported that adipose-specific mitochondrial dysregulation causes adipocyte energy deficiency and fat loss (Vernochet et al., 2014; Yang et al., 2016). Taken together, BA-induced adipocyte mitochondrial failure may result in reduced fat accumulation in the DKO mice.

We mapped this brown fat defect in the DKO mice to declined *Ucp1* expression and aging. UCP1 is responsible for brown adipose thermogenesis (Fedorenko et al., 2012), such that *Ucp1* deficient mice display mitochondrial disruptions, defective thermoregulation, and are resistant to diet-induced obesity with reduced white fat mass when housed at room temperature (Enerback et al., 1997; Kazak et al., 2017; Liu et al., 2003). Similarly, aged mice also show downregulation of UCP1 levels, mitochondrial dysfunction, reductions in brown fat heat production, and fat loss (Florez-Duquet and McDonald, 1998; Zoico et al., 2019). DKO mice and BA-induced adipocyte cultures exhibit overlapping phenotypes with both *Ucp1* knockout and aged mice (Table 3), suggesting that BA overload-induced mitochondrial dysfunction correlates to reducing *Ucp1* levels and brown adipose defects during aging.

**Table 3.**
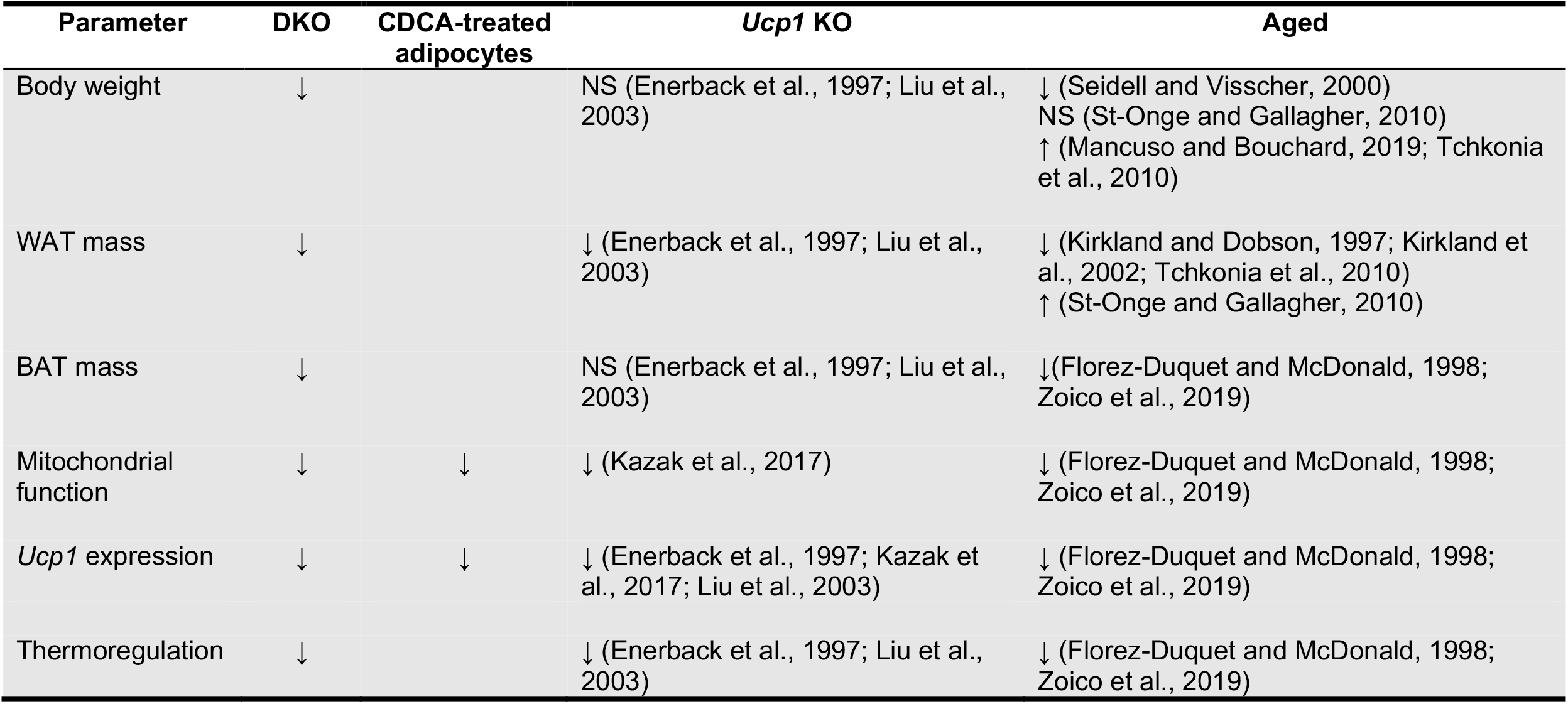
Comparison summary between primary adipocyte cultures upon CDCA treatment, DKO, *Ucp1* KO, and aged mice housed at room temperature.

Overall, our findings uncover a novel role for elevated levels of BAs in regulating brown adipose mitochondrial function and the subsequent thermogenesis.

## STAR METHODS

Detailed methods are provided in the online version of this paper and include the following:

- **KEY RESOURCES TABLE**
- **RESOURCE AVAILABILITY**
  ∘ Lead Contact
  ∘ Materials Availability
  ∘ Data and Code Availability
- **EXPERIMENTAL MODEL AND SUBJECT DETAILS**
  ∘ Human Adipose Tissue
  ∘ Animals
- **METHOD DETAILS**
  ∘ Histology
  ∘ Quantitative Real-Time PCR
  ∘ Analysis of the Degree of Lipid Unsaturation
  ∘ Mitochondrial Respiratory Enzyme Activity Assay
  ∘ Bile Acid Analysis
  ∘ Adipocyte Cell Culture
  ∘ Bile Acid Accumulation Assay
  ∘ Mitochondrial Membrane Potential Assay
  ∘ Cell Viability Assay
- **QUANTIFICATION AND STATISTICAL ANALYSIS**

## ACKNOWLEDGEMENTS

This work was supported by start-up funds from the University of Illinois at Urbana-Champaign (to SA) and R01 DK113080 from NIDDK (to SA). The authors are grateful to Ms. Oludemilade Akinrotimi for her assistance with obtaining transmission electron microscopy samples for analysis, Mr. Shawn D’Souza for qRT-PCR, and Dr. Lee-Jun Wong’s laboratory at Baylor College of Medicine for mitochondrial complex analysis. We also thank Ms. Lou Ann Miller from the Biological Electron Microscopy core at Materials Research Laboratory for performing transmission electron microscopy as well as the NIH West Coast Metabolomics Center at the University of California, Davis for performing adipose tissue BA analysis. We want to acknowledge the Microscopy Suite at the Beckman Institute, the University of Illinois at Urbana-Champaign for the access to the Raman micro-spectrometer used in the study.

## AUTHOR CONTRIBUTIONS

W.Z., P.V. and S.A. conceived and designed research; B.M.R., M.T.G. and M.B. supplied human adipose samples; W.Z., P.V., C.Z., R.R. and S.A. performed experiments; W.Z. analyzed data; W.Z. and S.A. interpreted data; W.Z. generated figures; W.Z. and S.A. drafted the manuscript; all authors were involved in editing and revising the manuscript, and had final approval of the submitted and published versions.

## STAR METHODS

### KEY RESOURCES TABLE

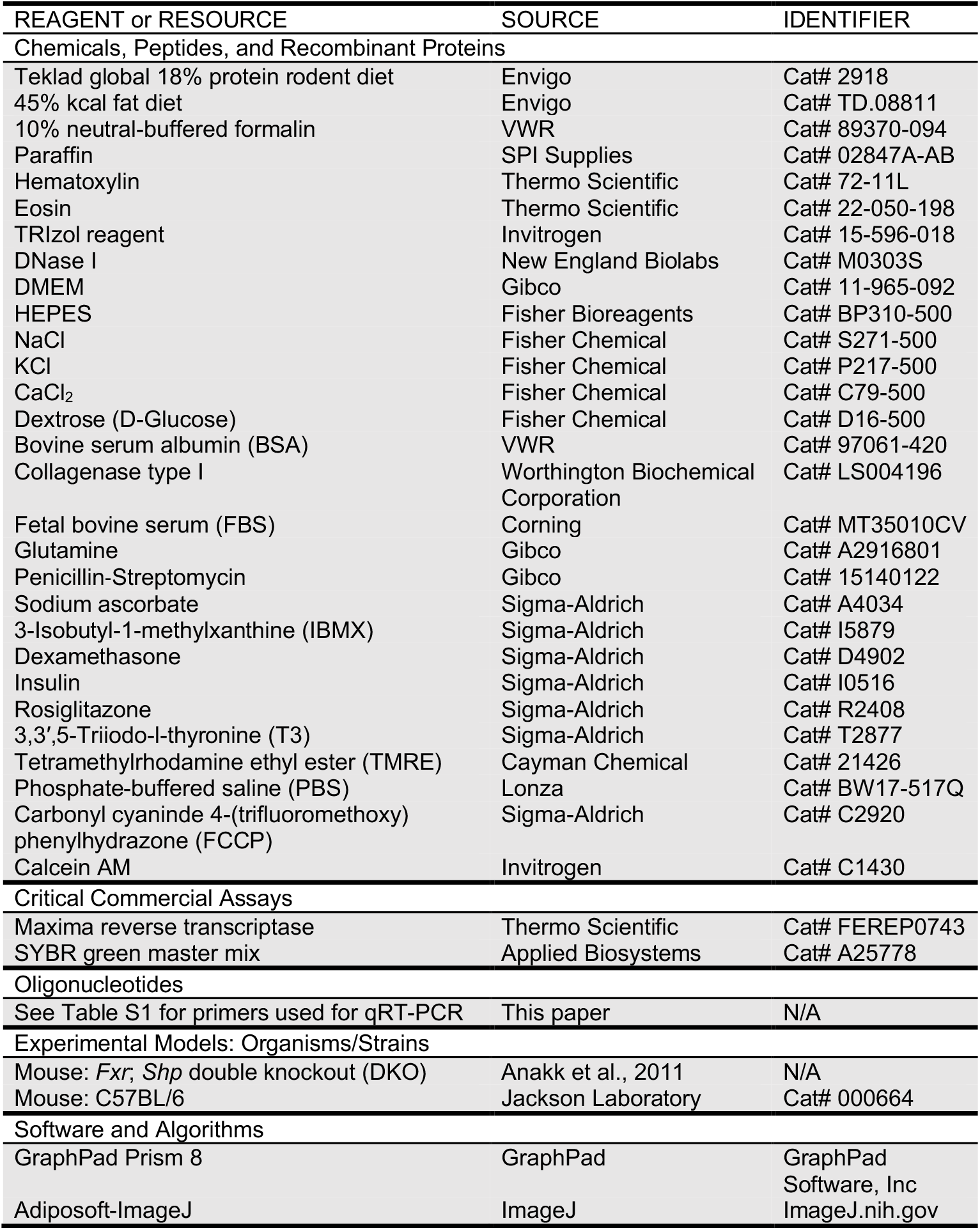

## RESOURCE AVAILABILITY

### Lead Contact

Further information and requests for resources and reagents should be directed to and will be fulfilled by the Lead Contact, Dr. Sayeepriyadarshini Anakk.

### Materials Availability

This study did not generate new unique reagents.

## Data and Code Availability

This study did not generate any unique datasets or code.

## EXPERIMENTAL MODEL AND SUBJECT DETAILS

### Human Adipose Tissue

Human perigastric white adipose tissue samples were obtained from tissue normally discarded from obese patients undergoing Roux-en-Y gastric bypass surgery (Robinson et al., 2019). All procedures were approved by the Institutional Review Board (IRB) at Carle Foundation Hospital and the University of Illinois at Urban-Champaign (IRB protocol # 14092). Informed consent was obtained from the subjects and the privacy rights of the subjects were observed.

### Animals

The generation of global Farnesoid X receptor (*Fxr*); Small heterodimer partner (*Shp*) double knockout (DKO) mice has been described (Anakk et al., 2011). Male DKO and C57BL/6 wild-type (WT) mice at 8- to 10-week-old were used. These mice were bred and maintained on a 12:12 light/dark cycle with *ad libitum* access to tap water and a normal chow diet in a climate-controlled (23 °C) animal facility at the University of Illinois at Urbana-Champaign. At 8-10 weeks, mice were fed a normal chow or 45% high-fat diet for 8 weeks to mimic obesogenic conditions and housed either at room temperature (RT) (23 °C) or at thermoneutrality (TN, 30 °C) to blunt brown fat thermogenic activity.

Mice were weighed weekly. After 8 weeks, a subset of the mice was used for monitoring core body temperature fluctuation using a Comprehensive Laboratory Animal Monitoring System (CLAMS). Briefly, animals were surgically implanted with a transmitter in the abdominal cavity and acclimated to the CLAMS cages. Fluctuations in the body temperature were recorded over the subsequent 24-hour period. The rest of mice were sacrificed at the end of the experimental regimen. Interscapular brown, inguinal and gonadal white adipose tissues were collected for primary preadipocyte culture and analysis of histology, gene expression, the degree of lipid unsaturation, mitochondrial respiratory enzyme activity, and bile acid levels. All experiments were performed following the National Institutes of Health guidelines for the care and use of laboratory animals, and all procedures were approved by the Institutional Animal Care and Use Committee at the University of Illinois at Urbana-Champaign.

## METHOD DETAILS

### Histology

Adipose tissues were fixed in 10% neutral-buffered formalin for 24 hours at 4 °C and processed. Formalin-fixed tissues were embedded using paraffin and cut on a microtome at 5 μm thickness and mounted onto charged glass microscope slides. Adipose sections were deparaffinized and stained with hematoxylin & eosin using standard histological protocol. Adipocyte size was quantified using Adiposoft-ImageJ software. Transmission electron microscopy (TEM) was performed as previously described (Nardi et al., 2019).

### Quantitative Real-Time PCR

Total RNA was isolated from primary pre- and post-differentiated adipocytes as well as snap-frozen adipose tissues using TRIzol reagent. Upon DNase I treatment, RNA was reverse transcribed into cDNA using a Maxima reverse transcriptase kit. Quantitative real-time PCR (qRT-PCR) was performed with SYBR green master mix using Applied Biosystems QuantStudio 7 Flex Real-Time PCR System. To determine relative expression values, the 2^−ΔΔCt^ method was used, where triplicate Ct values for each sample were averaged and subtracted from those derived from housekeeping gene *36B4*. All primers used are listed in Table S1.

### Analysis of the Degree of Lipid Unsaturation

Adipose tissues were fixed in 10% neutral-buffered formalin for 24 hours at 4 °C. Raman spectra of the adipose tissue were acquired using a Horiba LabRAM HR Raman spectroscopy imaging system. We used a 532 nm laser with an average power of 80 mW for excitation. The spectral acquisition speed was 5 s. A confocal configuration was applied using the spectrometer with a pinhole size of 200 μm. The Raman spectra background removal and intensity analysis were performed using Origin 2020. The relative degree of saturation level in formalin-fixed adipose tissues was measured using the Raman peak ratio 3006/1441 cm^−1^ (You et al., 2016).

### Mitochondrial Respiratory Enzyme Activity Assay

Activities of mitochondrial respiratory enzymes (i.e. complex I (NADH: ubiquinone oxidoreductase), complex III (ubiquinol-cytochrome c oxidoreductase), complex IV (cytochrome c oxidase)) were measured using a temperature-controlled spectrophotometer (Ultraspec 6300 pro, Biochrom Ltd.) as previously described (Brautbar et al., 2008). Citrate synthase was used as a marker for mitochondrial content. Enzyme activities are expressed as nmol/min/mg of protein.

### Bile Acid Analysis

Adipose bile acid (BA) analysis was performed in the NIH West Coast Metabolomics Center at the University of California, Davis. Adipose BAs were extracted from interscapular brown and gonadal white adipose tissue samples (4-4.75 mg) as previously described (La Frano et al., 2017). Six internal standards (GCA-d4, TCDCA-d4, CA-d6, GCDCA-d4, CDCA-d4, DCA-d4) were added. BA levels were quantified by ultra-high performance liquid chromatography chromatography-triple quadruple mass spectrometry (UHPLC-TQ-MS/MS) (Thermo Fisher Scientific). Data were processed with Metaboanalyst 4.0 (https://www.metaboanalyst.ca/MetaboAnalyst/home.xhtml) (Xia et al., 2009). The concentrations of individual BAs were summed to derive the levels of total, primary, secondary, unconjugated and taurine-conjugated BAs (Fu et al., 2012). BA hydrophobicity index was calculated as previously reported (Fu et al., 2012; Heuman, 1989).

### Adipocyte Cell Culture

Interscapular brown and inguinal white preadipocytes were isolated and cultured as described previously (Cannon and Nedergaard, 2001; Gao et al., 2017). Briefly, male C57BL/6 WT mice at 3- to 4-week-old were euthanized by isoflurane inhalational anesthesia followed by cervical dislocation. The interscapular brown and inguinal white adipose depots were harvested and minced with scissors in DMEM. Tissue fragments were incubated with digestion buffer ((ddH_2_O containing HEPES (100 mM), NaCl (123 mM), KCl (5 mM), CaCl_2_ (1.3 mM), glucose (5 mM), bovine serum albumin (BSA) (1.5% w/v) and collagenase type I (2 mg/mL)), and shaken at 300 rpm at 37 °C for 1 hour. The digested solution was then passed through a 100-μm cell strainer and placed on ice for 20 min. The infranatant below the top mature adipocyte layer was collected and passed through a 70-μm cell strainer, followed by centrifugation at 500 g for 10 min. The cell pellet was washed with DMEM, centrifuged, and then resuspended with pre-adipocyte culture medium (DMEM containing 10% fetal bovine serum (FBS), glutamine (4mM), HEPES (10 mM), Penicillin-Streptomycin (100 U/mL) and sodium ascorbate (25 μg/mL)). Two days post-confluency (Day 0), the brown or white preadipocytes were stimulated with preadipocyte induction medium (preadipocyte culture medium containing 3-Isobutyl-1- methylxanthine (IBMX) (0.5 mM), dexamethasone (1 μM), insulin (1 μg/mL) and rosiglitazone (1 μM) in the presence or absence of 3,3′,5-Triiodo-l-thyronine (T3) (1 nM)). Two days later (Day 2), the induction medium was changed to adipocyte maintain medium (preadipocyte culture medium containing insulin (1 μg/mL) with or without T3 (1 nM)). The maintain medium was changed every two days, and full differentiation of brown and white adipocytes was achieved by Day 6 and 8, respectively.

### Bile Acid Accumulation Assay

Post-differentiated adipocytes and Hep G2 cells (positive control) were treated with a fluorescein-labeled BA, choly-lysyl-fluorescein (CLF, 5 μM) for 24 hours at 37 °C, followed by nuclear counterstain with Hoechst 33342 for 20 min (Barber et al., 2015; Kostrubsky et al., 2003). Cell cultures were then washed three time with PBS and imaged using an EVOS fluorescent microscope.

### Mitochondrial Membrane Potential Assay

Mitochondrial membrane potential in live cells was determined by measuring the accumulation of tetramethylrhodamine ethyl ester (TMRE) in active mitochondria. After 24 hours BA treatment, post-differentiated adipocytes were stained with TMRE (250 nM) for 30 min at 37 °C, and then washed once with 0.2% BSA/PBS. The fluorescence of TMRE stained mitochondria was then measured using a microplate reader at Ex/Em = 544/590 nm. As a positive control for depolarization, uncoupler carbonyl cyaninde 4-(trifluoromethoxy) phenylhydrazone (FCCP) (20 μM) was applied to cells 10 min before TMRE labeling.

### Cell Viability Assay

Cell viability was measured using a cell-permeant dye, calcein AM. After BA treatment for 24 hours, post-differentiated adipocytes were stained with calcein AM (1 μM) for 30 min at 37 °C. The fluorescence in cell samples was then measured using a microplate reader at Ex/Em = 485/538 nm.

## QUANTIFICATION AND STATISTICAL ANALYSIS

Data were expressed as means ± SEM. Statistical analyses were performed using GraphPad Prism 8 software. Differences between two groups were analyzed using Student’s *t* test, and multiple group comparisons were analyzed using a one-way or two-way ANOVA with a Fisher’s LSD *post hoc* test. *P* < 0.05 was considered statistically significant.

