## Supplemental Figures and Tables for "Bile acid excess impairs thermogenic function in brown adipose tissue"

### Figure Legends

**Figure S1. DKO mice show altered adipocyte size and lipogenic gene expression profile upon normal chow diet**

(A) Distribution of adipocyte size of H&E-stained BAT (n=4-5 mice per group) and WAT (n=6-7 mice per group) sections of DKO and WT mice upon normal chow.

(B) Expression of genes related to lipogenesis in the BAT (n=7 mice per group) and WAT (n=5-7 mice per group) from DKO and WT mice upon normal chow.

(C) Representative Raman peak assignments of C=C and C-C bonds in the BAT and WAT from WT mice upon normal chow.

(D) The relative degree of lipid unsatuation (3006/1441 $cm^{-1}$) of BAT and WT from DKO and WT mice upon normal chow (n=4 mice per group).

Data are represented as mean ± SEM. *$P < 0.05$, **$P < 0.01$ compared to WT mice.

**Figure S2. Adipose mitochondrial gene transcript profile of WT and DKO mice upon normal chow diet**

(A-B) Expression of genes related to thermogenesis in the WAT (n=5-7 mice per group for RT, n=7-10 mice per group for TN) and BAT (n=7 mice per group for RT, n=7-10 mice per group for TN) from DKO and WT mice upon normal chow housed at RT or TN for 8 weeks.

Data are represented as mean ± SEM. *$P < 0.05$, **$P < 0.01$, ****$P < 0.0001$ compared to WT mice.

**Figure S3. Adipocyte size and lipogenic gene expression of DKO and WT mice upon chow at thermoneutral housing**

(A) Distribution of adipocyte size of H&E-stained BAT (n=7-11 mice per group, scale bar: 200 μm) and WAT (n=8-10 mice per group, scale bar: 200 μm) sections of DKO and WT mice upon normal chow housed at TN for 8 weeks.

(B) Expression of genes regulating lipogenesis in the BAT and WAT from DKO and WT mice housed at TN (n=7-10 mice per group).

Data are represented as mean ± SEM. *$P < 0.05$, **$P < 0.01$ compared to WT mice.

**Figure S4. DKO mice display decreased adipocyte size and reduced brown fat lipogenic gene expression upon HFD**

(A) Distribution of adipocyte size of H&E-stained BAT (n=5-9 mice per group for RT, n=5-6 mice per group for TN, scale bar: 200 μm) and WAT (n=6 mice per group for RT, n=5 mice per group for TN, scale bar: 200 μm) sections from WT and DKO mice housed at RT or TN upon HFD for 8 weeks.

(B-C) Expression of genes related to lipogenesis in the WAT (B) and BAT (C) from DKO and WT mice housed at RT or TN upon HFD (n=6-8 mice per group for RT, n=5-6 mice per group for TN).

Data are represented as mean ± SEM. **P* < 0.05, ***P* < 0.01, ****P* < 0.001, *****P* < 0.0001.

**Figure S5. DKO WAT exhibits altered levels of mitochondrial genes upon HFD**

Expression of genes regulating mitochondrial function in the WAT from WT and DKO mice housed at RT (n=6-8 mice per group) or TN (n=5-6 mice per group) upon HFD for 8 weeks.

Data are represented as mean ± SEM. **P* < 0.05, ***P* < 0.01, ****P* < 0.001, *****P* < 0.0001.

**Figure S6. High levels of CDCA do not affect adipocyte viability**

Cell viability of differentiated brown and white adipocytes upon CDCA treatment for 24 hours (n=4 cultures per group). Vehicle-treated adipocytes indicated by the dashed line.

Data are represented as mean ± SEM.

**Figure S7. CDCA alters brown adipocyte mitochondrial membrane potential in a dose-dependent manner**

Quantification of mitochondrial membrane potential of differentiated brown adipocytes upon different concentrations of CDCA treatment for 24 hours (n=4 cultures per group).

Data are represented as mean ± SEM. ***P* < 0.01, ****P* < 0.001, *****P* < 0.0001 compared to vehicle-treated adipocytes.

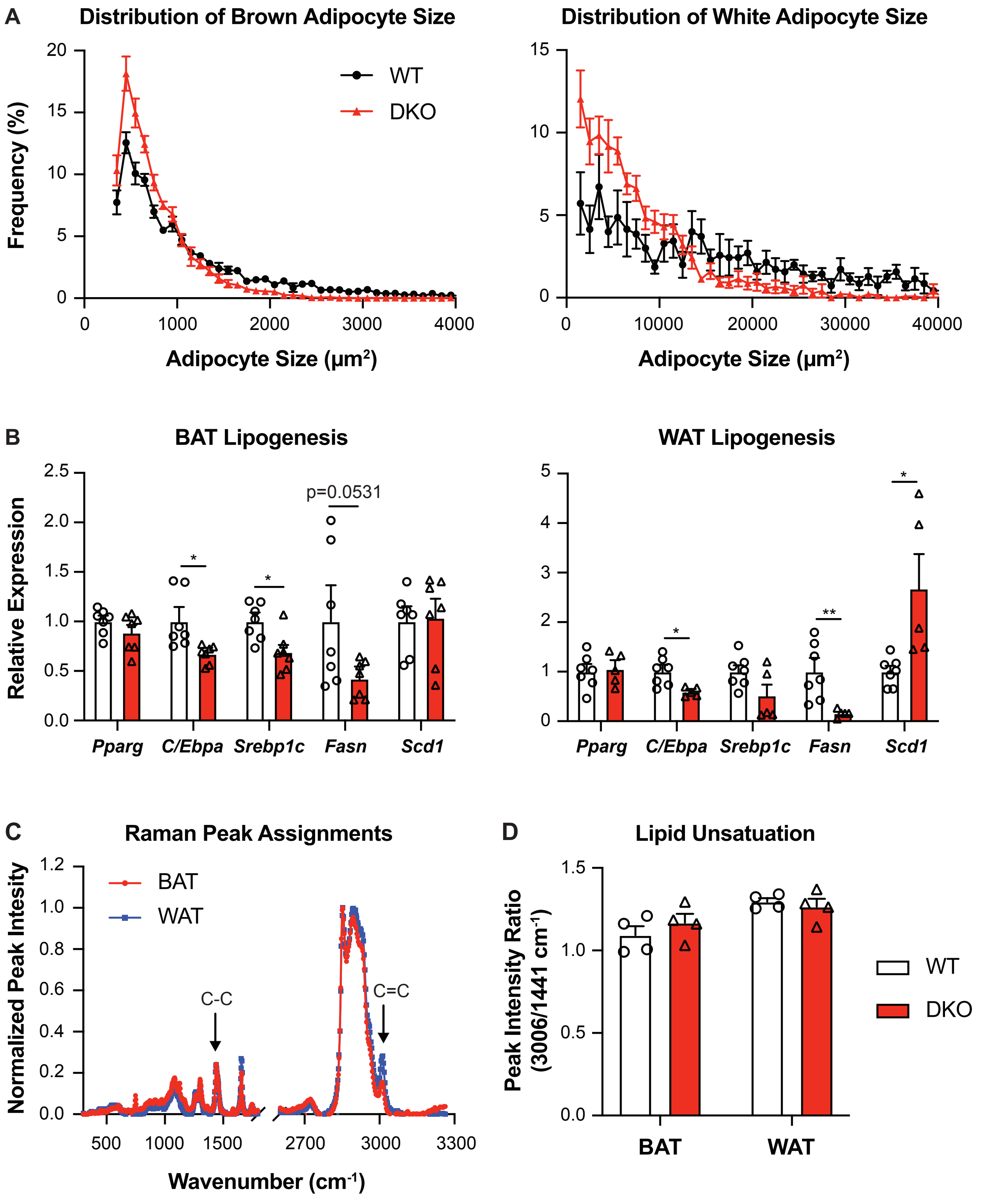
A
Distribution of Brown Adipocyte Size
Distribution of White Adipocyte Size
Frequency (%)
WT
DKO
Adipocyte Size (μm²)
Adipocyte Size (μm²)
B
BAT Lipogenesis
WAT Lipogenesis
Relative Expression
p=0.0531
Pparg
C/Ebpa
Srebp1c
Fasn
Scd1
C
Raman Peak Assignments
Normalized Peak Intesity
BAT
WAT
C-C
C=C
Wavenumber (cm-1)
D
Lipid Unsatuation
Peak Intensity Ratio (3006/1441 cm-1)
BAT
WAT
WT
DKO

Figure S1

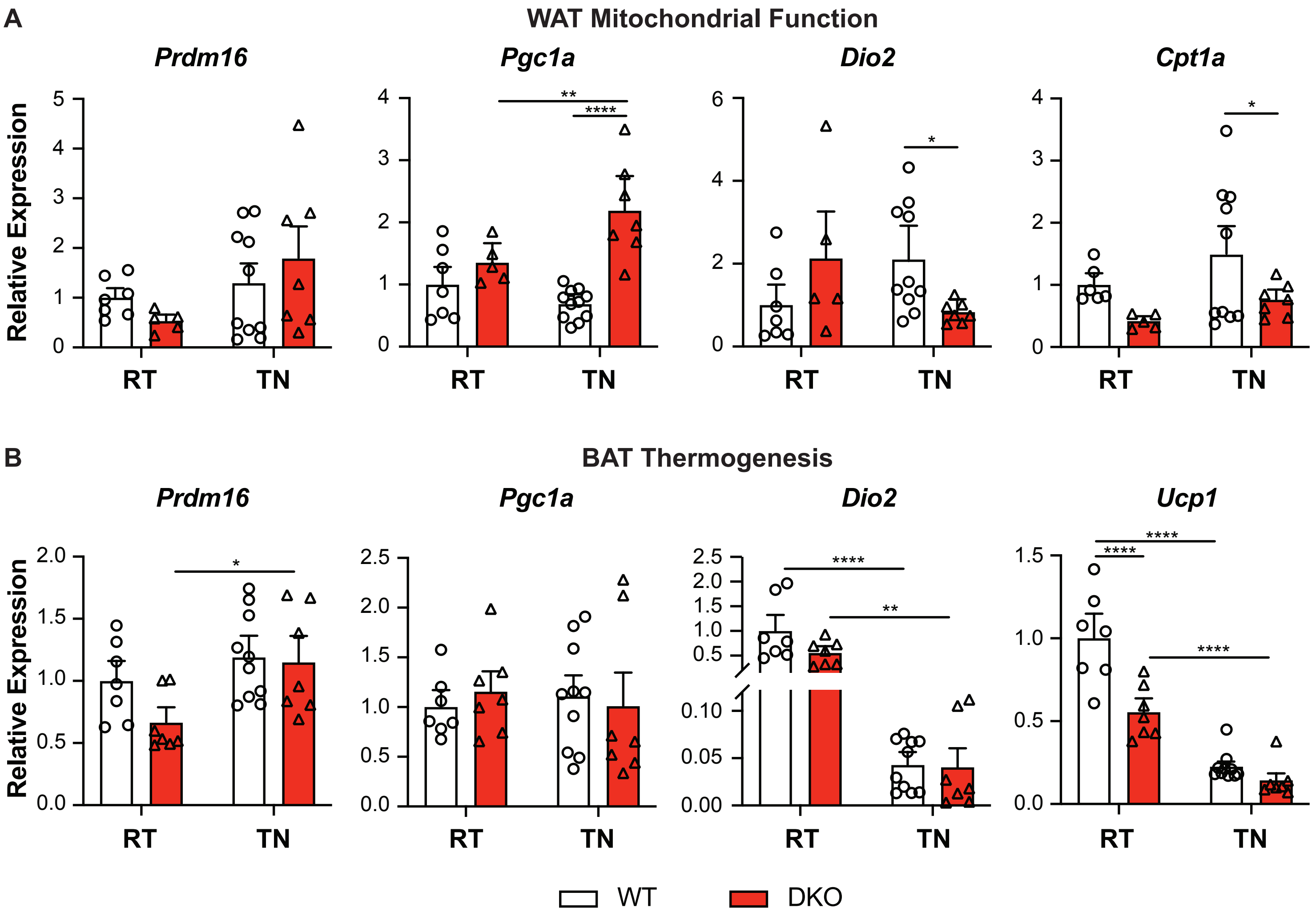
A
WAT Mitochondrial Function
Prdm16
Pgc1a
Dio2
Cpt1a
Relative Expression
RT
TN
B
BAT Thermogenesis
Prdm16
Pgc1a
Dio2
Ucp1
Relative Expression
RT
TN
WT
DKO

**Figure S2**

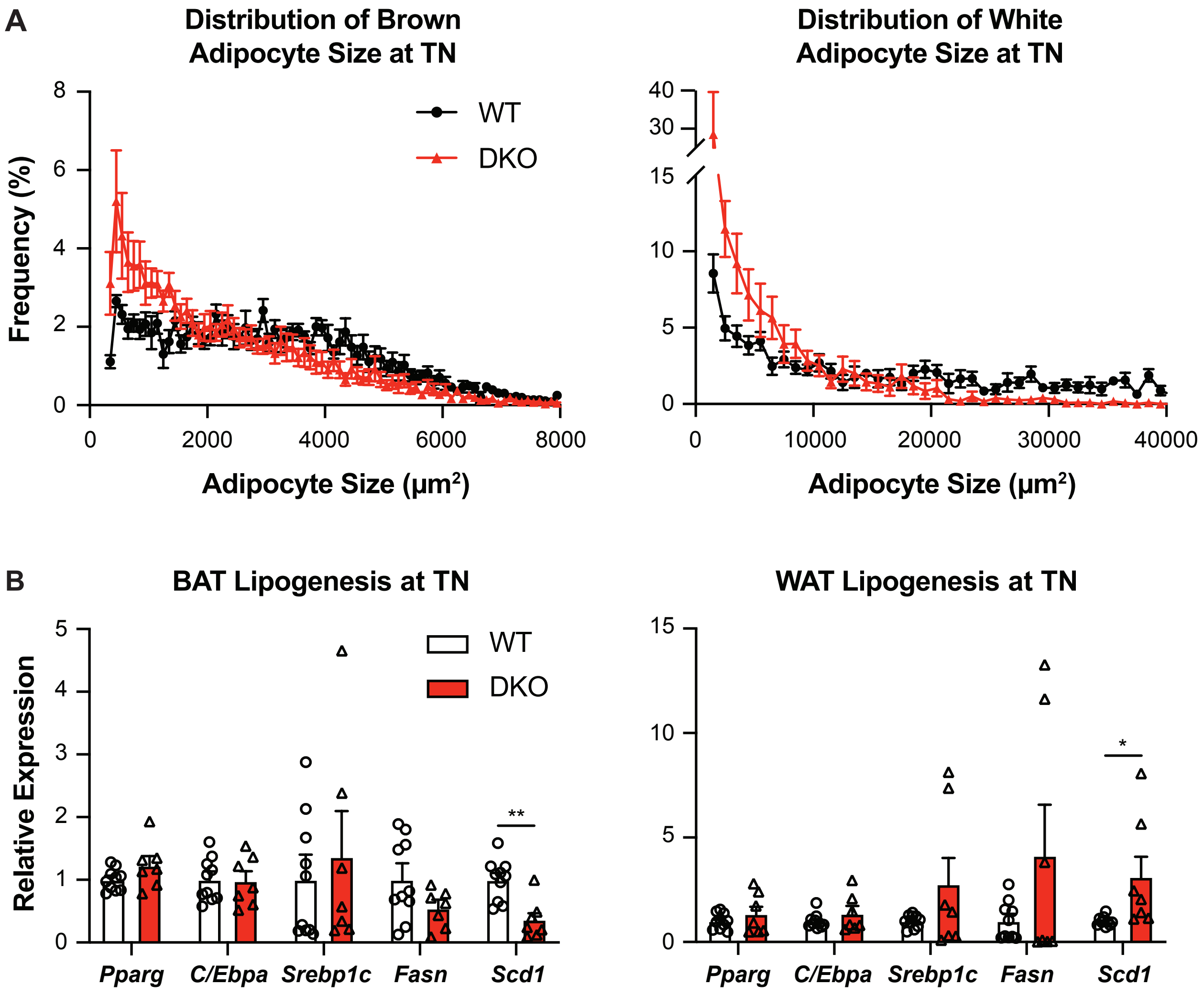
A
Distribution of Brown Adipocyte Size at TN
Frequency (%)
WT
DKO
Adipocyte Size (μm²)
Distribution of White Adipocyte Size at TN
Adipocyte Size (μm²)
B
BAT Lipogenesis at TN
Relative Expression
WT
DKO
**
Pparg
C/Ebpa
Srebp1c
Fasn
Scd1
WAT Lipogenesis at TN
*
Pparg
C/Ebpa
Srebp1c
Fasn
Scd1

**Figure S3**

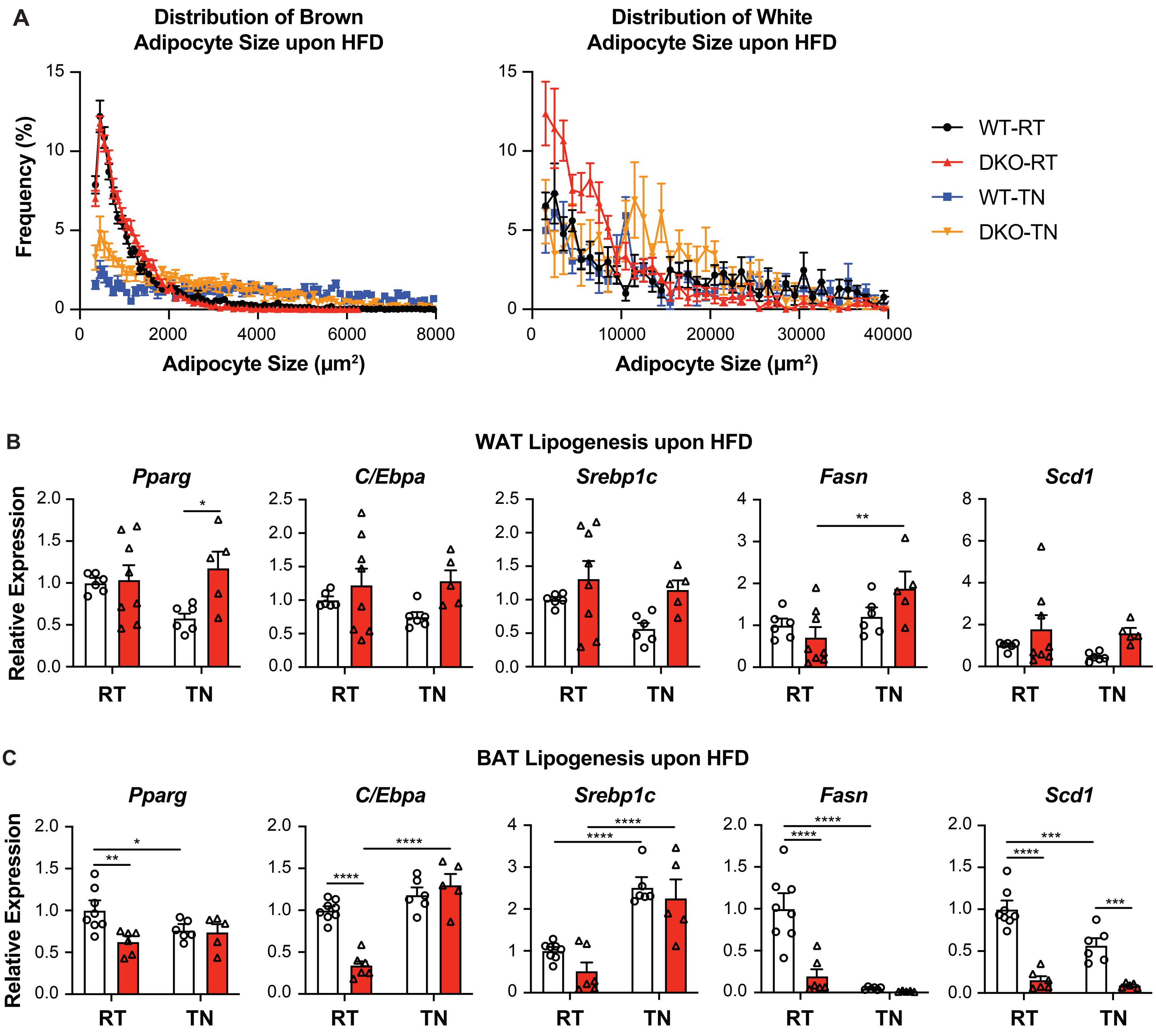

A
Distribution of Brown Adipocyte Size upon HFD
Distribution of White Adipocyte Size upon HFD
Frequency (%)
Adipocyte Size (μm²)
WT-RT
DKO-RT
WT-TN
DKO-TN
B
WAT Lipogenesis upon HFD
Pparg
C/Ebpa
Srebp1c
Fasn
Scd1
Relative Expression
RT
TN
C
BAT Lipogenesis upon HFD
Pparg
C/Ebpa
Srebp1c
Fasn
Scd1
Relative Expression
RT
TN


Figure S4

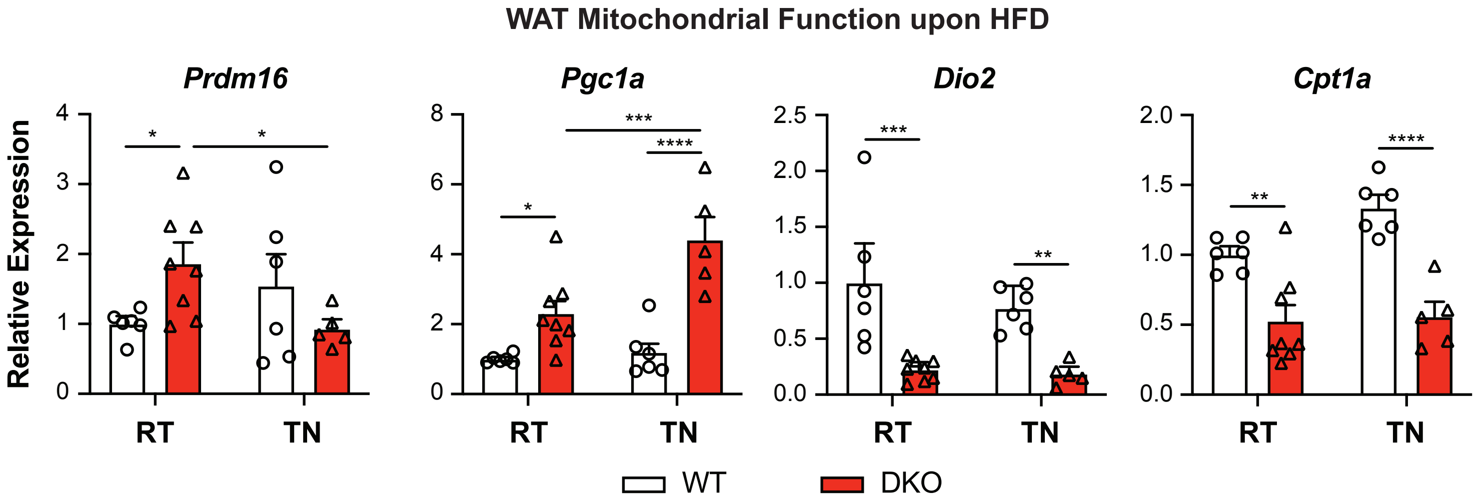
WAT Mitochondrial Function upon HFD
Prdm16
Pgc1a
Dio2
Cpt1a
Relative Expression
RT
TN
WT
DKO
*
**
***
****

**Figure S5**

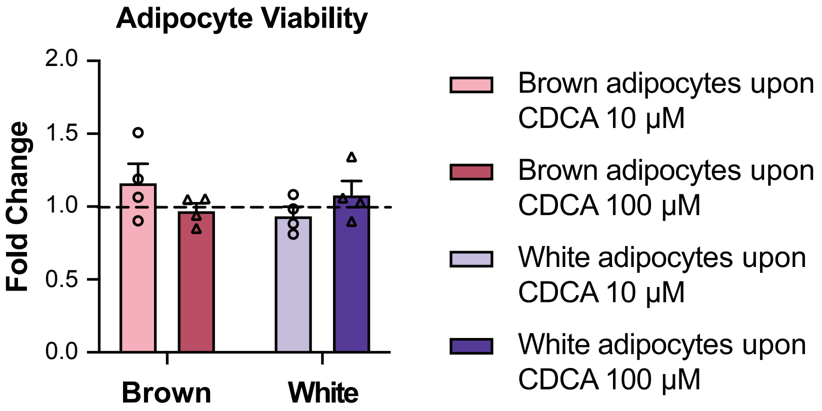
Adipocyte Viability
Fold Change
2.0
1.5
1.0
0.5
0.0
Brown
White
Brown adipocytes upon CDCA 10 μM
Brown adipocytes upon CDCA 100 μM
White adipocytes upon CDCA 10 μM
White adipocytes upon CDCA 100 μM

**Figure S6**

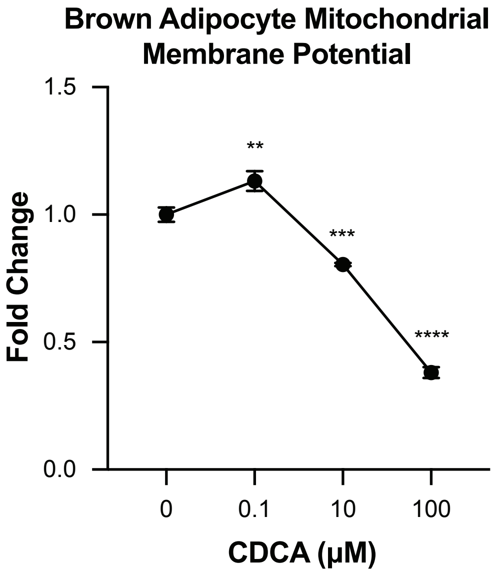
Brown Adipocyte Mitochondrial Membrane Potential
Fold Change
1.5
1.0
0.5
0.0
**
***
****
0
0.1
10
100
CDCA (μM)

**Figure S7**

| Gene | Forward (5' -> 3') | Reverse (5' -> 3') |
|---|---|---|
| *36B4 (mouse)* | AGATGCAGCAGATCCGCAT | GTTCTTGCCCATCAGCACC |
| *Prdm16* | GCCATGTGTCAGATCAACGA | CCTTCTTTCACATGCACCAA |
| *Pgc1a* | CCCACAGAAAACAGGAACAG | CTGGGGTCAGAGGAAGAGAT |
| *Dio2* | CAGTGTGGTGCACGTCTCCAATC | TGAACCAAAGTTGACCACCAG |
| *Ucp1* | GGCCTCTACGACTCAGTCCA | TAAGCCGGCTGAGATCTTGT' |
| *Cpt1a* | TGATGACGGCTATGGTGTTTC | CAAACAAGGTGATAATGTCCATC |
| *Pparg* | CAAGAATACCAAAGTGCGATCAA | GAGCTGGGTCTTTTCAGAATAATAAG |
| *C/Ebpa* | GCGGGAACGCAACAACATC | GTCACTGGTCAACTCCAGCAC |
| *Srebp1c* | GGAGCCATGGATTGCACATT | GGCCCGGGAAGTCACTGT |
| *Fasn* | GCTGCGGAAACTTCAGGAAAT | AGAGACGTGTCACTCCTGGACTT |
| *Scd1* | CCCTGCGGATCTTCCTTATC | TGTGTTTCTGAGAACTTGTGGTG |
| *Cyp7a1* | CCTAAGAGCAAAGCAAAGGAAAC | CTTTGTGGTATGACAGGGAGTT |
| *Cyp7b1* | GACGATCCTGAAATAGGAGCACA | AATGGTGTTTGCTAGAGAGGCC |
| *Cyp8b1* | TTTCTGAGGGAGCAAGGAATAG | GGAATAAGAGGACCCAGAAACA |
| *Cyp27a1* | CCTACATCCATTCGGCTCT | CCAGGGCAATCTCATACTTC |
| *Cyp2c70* | TGGCTTTCTCAGCAGGAAGAA | AACTGGCTTGGTGTCGATGT |
| *Slc27a5* | TACTGGAGAGGTGGAGTGTGT | CAACCTTACCCTCACACCCTG |
| *Baat* | GTGCTGGTGGATTGATGGAGT | CCGAGGACCTTAGGATGTCTC |
| *Slco1a6* | GATTGGTGTTGGTTGTGCAG | TGGGATCTGTTTTCCACACA |
| *Slco1b2* | TTCACCACAACAATGGCCTA | TTTTCCCCACAGACAGGTTC |
| *Slc51a* | TTGTGATCAACCGATTTGT | CCACGCCTGTTTCATTACCT |
| *Slc51b* | ATCCTGGCAAACAGAAATCG | GGGTCTGGCAGAAAGACAAG |
| *p16INK4a* | CCCAACGCCCCGAACT | GCAGAAGAGCTGCTACGTGAA |
| *p19ARF* | GCCGCACCGGAATCCT | TTGAGCAGAAGAGCTGCTACGT |
| *p21* | AGATCCACAGCGATATCCAGAC | ACCGAAGAGACAACGGCACACT |
| *IL6* | TCTATACCACTTCACAAGTCGGA | GAATTGCCATTGCACAACTCTTT |
| *36B4 (human)* | TCCTTTGGGCTGGTCATC | GCAGACAGACACTGGCAACA |
| *CYP7A1* | TGGCATCCTTCCCTTTCTAATC | TCCAAGAATAAGCCATAGACAACA |
| *CYP7B1* | GGCCCTCTGCTTGCTTGTC | GAAGCCAACCTTTTATCAATGGA |
| *CYP8B1* | CACCCGGCTAGTTTCTGTATT | AGTGGCTCATGCCTGTAATC |
| *CYP27A1* | CAACGGAGCTTAGAGGAGATTC | TGCAGTTGCAGGGCATAG |

**Table S1. Primer sequences used in qRT-PCR analysis**
